# Allosteric feed-forward activation of AP-2 *via* FCHO-BMP2K axis promotes clathrin-mediated endocytosis

**DOI:** 10.1101/870899

**Authors:** Shikha T. Ramesh, Kolaparamba V. Navyasree, Anjitha B. Ashok, Nishada Qathoon, Suryasikha Mohanty, Rajeeb K. Swain, Perunthottathu K. Umasankar

## Abstract

Spatio-temporal regulation of central adaptor complex, AP-2 is pivotal for clathrin-mediated endocytosis (CME). We recently discovered that FCHO proteins trigger clathrin-coated pit (CCP) formation by allosterically activating AP-2 on plasma membrane (Umasankar *et al*., 2014). Here, we demonstrate that this activation promotes AP-2 phosphorylation *via* recruitment and stabilization of BMP-2 inducible kinase (BMP2K), a *bona fide* AP-2 kinase leading to CCP maturation. Accordingly, BMP2K mislocalizes and degrades in FCHO knockout/ AP-2 depleted cells. Functional inactivation of kinase impairs AP-2 phosphorylation leading to altered lattice morphology and CME phenotypes reminiscent of CCP maturation defects. Reexpression of FCHO rescues AP-2 phosphorylation defects in FCHO knockout cells implying membrane activation of AP-2 is a prerequisite for kinase function. Furthermore, gain- and loss-of function phenotypes of FCHO and BMP2K are analogous and mirror altered AP-2 functions during zebrafish embryogenesis. Together, our findings reveal an *in vivo* allosteric feed-forward axis for operation of CME.

## Introduction

Clathrin-mediated endocytosis (CME) is a major vesicular transport mechanism in eukaryotes for internalization of cell surface transmembrane protein cargo (Traub, 2009). The process involves sequential transition of a flat piece of membrane into a cargo-laden pit (CCP) decorated with a cytosolic coat protein, clathrin (Merrifield and Kaksonen, 2014, Mettlen et al., 2018). CCP ultimately buds into cell interior with the help of dynamin GTPase, to become clathrin-coated vesicle (CCV) that after uncoating fuses with endosomes to deliver cargo (Schmid and Frolov, 2011; Ferguson and De Camilli, 2012). CCV biogenesis occurs in ∼60 seconds and is efficiently choreographed through temporally co-ordinated recruitment of adaptors and accessory proteins on the membrane (Taylor et al., 2011; Kaksonen and Roux, 2018). The incapability of clathrin to bind membrane and cargo necessitates evolution of a special set of adaptors generally termed clathrin-associated sorting proteins (CLASPs). CLASPs translocate from cytosol to the inner leaflet of plasma membrane to sort cargo, recruit clathrin and accessory proteins for CCP assembly (Traub and Bonifacino, 2013).

Adaptor protein complex-2 (AP-2) is a major heterotetrameric CLASP that consists of α, β2, µ2 and σ2 subunits (Robinson, 2004). The trunk domains of α and β2 together with the entire µ2 and σ2 form AP-2 core (Kirchhausen et al., 2014). The extended domains of α and β2 subunits form appendages that connect to the core through long flexible hinges (Traub et al., 1999; Owen et al., 2004). AP-2 facilitates two major functions in CME. First, through its core, AP-2 mediates binding to plasma membrane phospholipids (PIP2) and to sorting signals on cargo. AP-2 sorts cargo containing tyrosine based (YXXØ) and di-leucine based [DE]XXXL[LI] sorting motifs through µ2 and σ2subunits respectively (Traub, 2009). Second, AP-2 recruits clathrin and late stage accessory proteins through hinge and appendages. Together it promotes CCP initiation (Cocucci et al., 2012; Godlee and Kaksonen, 2013), assembly and maturation (Shih et al., 1995; Edeling et al., 2006). Thus, AP-2 efficiently orchestrates CCV biogenesis by co-ordinating multiple low-affinity interactions with binding partners (Schmid and Macmahon, 2007).

AP-2 undergoes series of conformational changes that functionally translate into distinct stages in CCV biogenesis (Kadlecova et al., 2017). Crystal structures suggest at least three functionally distinct conformations of AP-2; (1) closed, inactive state in which binding surfaces for PIP2, cargo and clathrin are masked (Collins et al., 2002), (2) open, active state in which these binding surfaces are exposed and aligned in a co-planar fashion (Kelly et al., 2008; Jackson et al., 2010; Kelly et al., 2014), and (3) a new open+ active conformation in which an additional threonine (T156) residue on the flexible linker region of µ2 is exposed for phosphorylation (Wrobel et al., 2019). Accordingly, large-scale mutagenesis screens in the nematode, *C.elegans* identify phenotypic functional mutations reflecting different conformations of AP-2 (Hollopeter et al., 2014; Beacham et al., 2018, Partlow et al., 2019).

Several allosteric regulators have been proposed in mediating AP-2 conformational transitions. *In vitro* structural and biochemical studies supported by quantitative live-cell imaging analyses suggest PIP2 and cargo binding promotes transition from closed to open state stabilizing AP-2 on membrane (Honing et al., 2005; Jackson et al., 2010; Kadlecova et al., 2017). Recent investigations establish role of muniscin family of proteins (FCHO1, FCHO2 and neuronal SGIP1) (Reider et al., 2009; Henne et al., 2010; Umasankar et al., 2012) in this process. In genome-edited HeLa cells lacking FCHO (Umasankar et al., 2014) and in *C. elegans fcho* mutants (Hollopeter et al., 2014), majority of AP-2 fails to recruit on membrane, hampering CCP inception. Residual CME still persists in these cells (Mulkearns et al., 2012, Umasankar et al., 2014) by virtue of additional AP-2 interactions mediated by coat-initiating proteins, EPS15/R (Ma et al., 2016). Consistently; nanobody-mediated disruption of EPS15/R-AP-2 interaction further deteriorates CME in 1.E cells by completely inactivating AP-2 (Traub, 2019) suggesting parallel mechanisms for AP-2 activation. In all these cases, actual *in vivo* conformation of AP-2 remains unclear. Nevertheless, expression of intrinsically disordered linker region of muniscins that binds AP-2 core restores AP-2 function in 1.E cells and in *C. elegans* (Umasankar et al., 2014; Hollopeter et al., 2014) confirming allosteric activation. More recent studies demonstrate role of another endocytic protein, NECAP in allosteric regulation of AP-2 (Ritter et al., 2013; Beacham et al., 2018). It has been shown that in mammalian cells, NECAP promotes AP-2 activation by coupling open+ phosphorylated forms to downstream effectors of CCV biogenesis (Wrobel et al., 2019). On the contrary, studies in *C. elegans* suggest that NECAP inactivates AP-2 by converting open to closed state (Beacham et al., 2018; Partlow et al., 2019).

Reversible phosphorylation of T156 µ has been proposed an important allosteric regulator of AP-2 function (Wild and Brodsky, 1996; Smythe, 2002). However, several aspects of AP-2 phosphorylation remain unclear and have been largely debated. For instance, when AP-2 phosphorylation occurs during CCV biogenesis and its functional significance remain poorly understood. Some studies indicate phosphorylation associates with open AP-2 based on the phenotypes in *C. elegans* and on the structural predictions of T156 µ linker positioning (Hollopeter et al., 2014; Beacham et al., 2018; Partlow et al., 2019). It is also disputed whether phosphorylation opens AP-2 (Höning et al., 2005; Jackson et al., 2010), closes AP-2 (Beacham et al., 2018; Partlow et al., 2019) or induces an entirely different conformation (Wrobel et al., 2019). In addition, earlier *in vitro* structural and biochemical studies predict AP-2 phosphorylation may increase cargo binding affinity (Pauloin and Thurieau, 1993; Olusanya et al., 2001; Ricotta et al., 2002).

Recent structural and live-cell imaging approaches attempt to bring in more clarity to this picture (Kadlecova et al., 2017; Wrobel et al., 2019; Partlow et al., 2019). New crystal structures of phosphorylated AP-2 clearly indicate that T156 µ site is not accessible in open state, thus cannot be phosphorylated by kinases. However, it is predicted that active kinase can phosphorylate open+ or closed states and has access to AP2 where those conformations exist (Wrobel et al., 2019). These studies also show that phosphorylated AP-2 creates new sites for NECAP engagement without perturbing cargo and clathrin binding to either couple CCP for maturation (Wrobel et al., 2019) or to inactivate and recycle AP-2 (Partlow et al., 2019). Additionally, these studies find T156 μ phosphorylates early at CCP formation and increases as CCP grows suggesting a positive role for AP-2 phosphorylation (Kadlecova et al., 2017; Wrobel et al., 2019).

Incongruity in AP-2 phosphorylation studies may arise largely due to lack of understanding of specific kinases that phosphorylate AP-2 and their regulatory mechanisms in cells and organisms. Earlier investigations help identify Adaptor associated kinase-1 (AAK1) that phosphorylates AP-2 µ2 in *in vitro* assays (Conner and Schmid, 2002). Accordingly, AAK1 co-purifies with AP-2 in CCV, binds to AP-2/ clathrin and localizes to CCPs in cells (Conner et al., 2003; Jackson et al., 2003). Yet, RNAi-mediated silencing of AAK1 fails to induce any significant changes in AP-2 phosphorylation status or cargo uptake in cells (Conner and Schmid, 2003; Henderson and Conner, 2007). This leaves the possibility of potential redundant kinases and novel regulatory mechanisms for AP-2 phosphorylation.

BMP2K was originally identified as a serine-threonine kinase regulating osteoblast differentiation whose gene product is sharply up-regulated by BMP-2 stimulation (Kearns et al., 2001). Later, various genetic and proteomic screens link BMP2K to HIV entry (Zhou et al., 2008), developmental dysplasia of the hip (Zhao et al., 2017), myopia (Liu et al., 2009), leukemia (Brehme et al., 2009; Pandzic et al., 2016), breast cancer metastasis (Mercado-Matos et al., 2018) *etc*. Recent multivariate proteomic profiling approaches identify BMP2K as a component of purified CCV from HeLa cells (Borner et al., 2012). Moreover, structural similarities of kinase domains (75% identical) and *in vitro* inhibition profiles categorize BMP2K as a member of Numb-associated protein kinase (NAK) family and as the closest homolog of AAK1 (Krieger et al., 2013; Elkins et al., 2016; Sorrell et al., 2016).

Here, by integrating reverse genetic, pharmacological and biochemical approaches in genome-edited cells and *D. rerio* (zebrafish) embryos, we unravel BMP2K as a *bona fide* AP-2 kinase that reversibly phosphorylates µ2 at T156 in a spatio-temporally controlled fashion to regulate AP-2 functions. We demonstrate that membrane activated AP-2 induced by FCHO triggers recruitment and stabilization of BMP2K at CCPs. Accordingly, FCHO knockouts defective in membrane activation of AP-2 or BMP2K mutants that cannot localize to CCPs fail to induce AP-2 phosphorylation. Inhibition of BMP2K activity in HeLa cells abolishes AP-2 phosphorylation and elicits CME phenotypes temporally distinct from FCHO knockouts. In *D. rerio* embryos, gain- and loss-of-function of FCHO and BMP2K perturb AP-2 functions leading to aberrant CME during development. Together, our findings propose a feed-forward allosteric axis involving FCHO-BMP2K that activates AP-2 for proper functioning of CME.

## Results

### Loss of FCHO identify BMP2K as a *bona fide* AP-2 kinase

Previously, we had identified CME defects in 1.E (FCHO knockout) cells occurring due to altered distribution and function of AP-2 in the absence of FCHO. Immunofluorescence with the AP.6 monoclonal antibody confirms the altered distribution of AP-2 positive structures in 1.E cells (**Figure 1A and B**, quantified in **Figure 7C**). A 5 minute pulse of fluorescent transferrin followed by acid stripping of surface ligands (endocytosis assay-protocol #2) shows reduced fluorophore in peripheral endosome compartments in 1.E cells validating endocytic defects (**Figure 1A’, A”, B’ and B”**). To investigate if membrane activation of AP-2 is required for phosphorylation, we compared phosphorylation status of AP-2 in 1.E cells and parental HeLa SS6 cells. A commercial antibody specific to phosphorylated T156 µ (pT156 µ) reveals the extent of AP-2 phosphorylation in HeLa cells. On blots, steady-state levels of pT156 µ dramatically reduce in 1.E as compared to HeLa SS6 (**Figure 1C**), indicating defective µ2 phosphorylation in the absence of FCHO. Reduction in phosphorylation is not due to destabilization and degradation of AP-2 complex as steady state levels of µ2 and other AP-2 subunits are similar in 1.E and HeLa SS6 (**Figure 1C**). Given that µ2 phosphorylation is driven by NAKs, we checked if phosphorylation defect is due to loss of AAK1 or BMP2K in 1.E cells. Though both these kinases are invariably present in low copy numbers across cell types, BMP2K is reported to be slightly more abundant (Hein et al., 2015) and more membrane associated (Itzhak et al., 2016) than AAK1 in HeLa cells. Consistently, immunoblotting with different commercially available antibodies against both these kinases could detect only BMP2K in HeLa lysate in our experiments. We find that steady state levels of BMP2K drastically decline in 1.E cells compared to HeLa SS6 with unchanged levels of AP-2, clathrin and other clathrin-associated adaptor and accessory proteins (**Figure 1D**).

**Figure 1:**
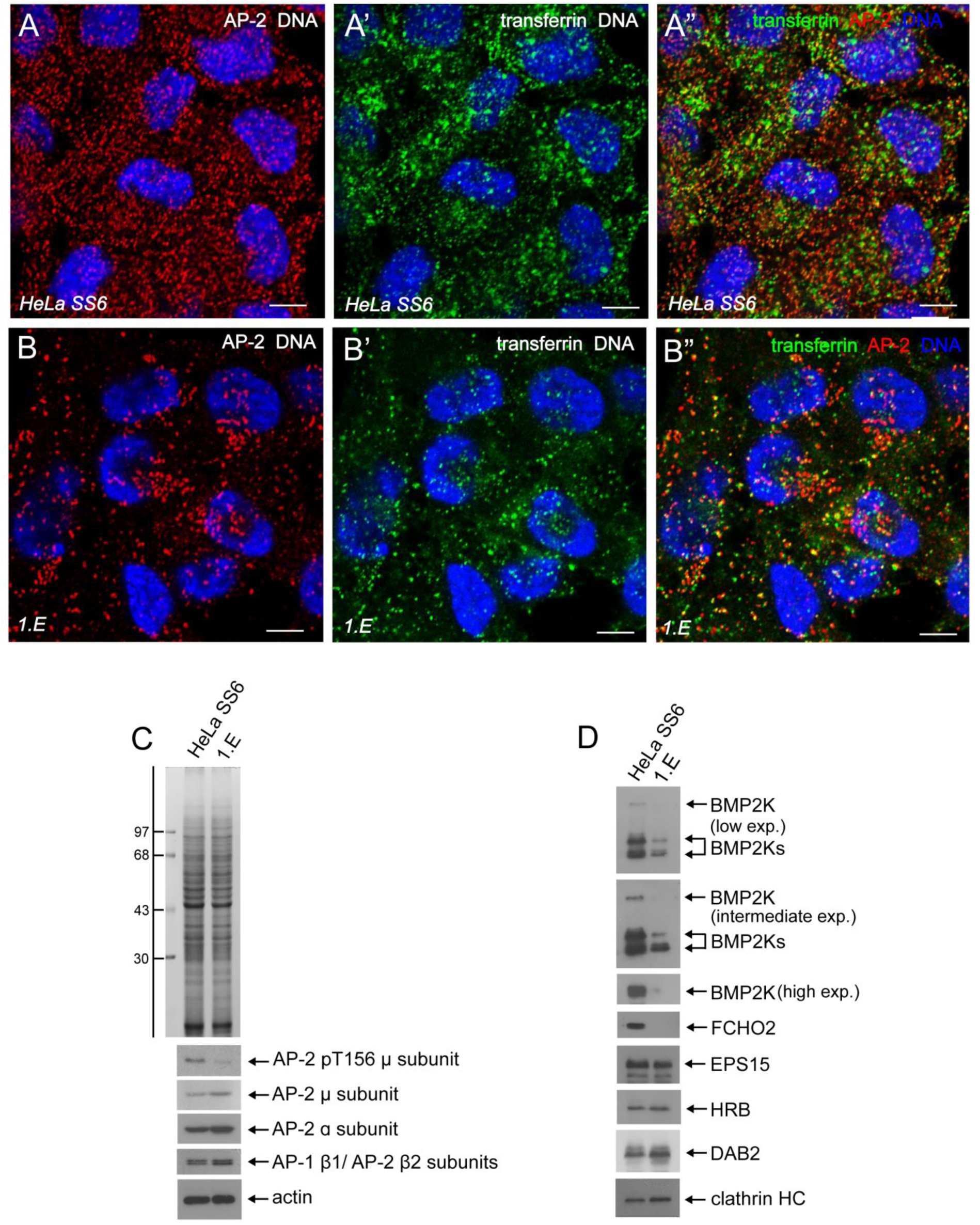
AP-2 phosphorylation defects in 1.E cells lacking FCHO1/2. **(A and B)** Representative confocal optical sections of the basal plane of parental HeLa SS6 cells (**A-A”**) and 1.E cells (**B-B”**) pulsed with Alexafluor 488-transferrin (green) for 5 minutes followed by acid stripping (protocol #2) and immunolabeling with anti-AP-2 α subunit mAb AP.6 (red) and Hoechst DNA dye (blue) as indicated. Images with all three colours merged are shown in **A” and B”**. Scale bar-10 µm. **(C and D)** SDS-PAGE and immunoblot analysis of whole cell lysates from HeLa SS6 and 1.E. Gels were either stained with Coomassie blue (upper panel in C) or transferred to nitrocellulose and probed with the indicated antibodies. Low, intermediate and high exposures of autoradiographic films detecting large (BMP2K) and small (BMP2Ks) isoforms of kinase are indicated. Molecular mass standards (in kDa) are shown on the left.

Since reduced phosphorylation correlates with loss of BMP2K, we examined if BMP2K is a *bona fide* AP-2 kinase in cells. Quantitative immunofluorescence analyses with pT156 µ and AP.6 antibodies reveal basal mean fluorescent intensity ratios of phosphorylated AP-2 to total AP-2 in majority of CCPs in HeLa SS6 cells (**Figure 2A, B and Figure 2- figure supplement 1**). Transient expression of a C-terminally GFP-tagged BMP2K (BMP2K-GFP) populates AP-2 positive CCP on the ventral surface of HeLa SS6 cells (**Figure 2A**). Strikingly, mean fluorescent intensity ratio of pT156 µ/ AP-2 increases ∼ 3 fold in BMP2K-GFP positive CCP (**Figure 2A, B and Figure 2- figure supplement 1**) suggesting enhanced AP-2 phosphorylation by excess kinase. GFP-tagging at the C-terminus causes negligible effect on kinase function. Immunoblots show comparable increase in the steady state levels of pT156 µ in cells transfected with GFP-tagged and- untagged versions of BMP2K (**Figure 2C**). In addition, AP-2 phosphorylation status in BMP2K transfected cells is comparable to that of cells treated with calyculin, a pan-serine/threonine phosphatase inhibitor that reversibly arrests proteins in phosphorylated state (**Figure 2C**). Recent pharmacological studies identify SGC-AAK1-1 (Elkins et al., 2016; Agajanian et al., 2019), a small molecule functionally similar to the earlier version of brain-penetrant inhibitor LP-935509 (Wrobel et al., 2019) that potently and selectively inhibits NAKs both *in vitro* and *in vivo*. Co-crystal structures of inhibitor-bound kinase confirm that inhibition is mediated through competitive binding to the conserved ATP site on the kinase domain (Agajanian et al., 2019). We used SGC-AAK1-1 to inhibit AP-2 kinase (AAK1 and BMP2K) functions in HeLa cells. Steady state levels of pT156 µ decrease on blots with increasing concentrations of SGC-AAK1-1 confirming specificity and efficiency of the inhibitor (**Figure 2D**). The inhibitor also suppresses elevated levels of pT156 µ in BMP2K expressing cells suggesting AP-2 phosphorylation impedes upon BMP2K inactivation (**Figure 2E**). Together, these results establish BMP2K a valid AP-2 kinase in cells.

**Figure 2:**
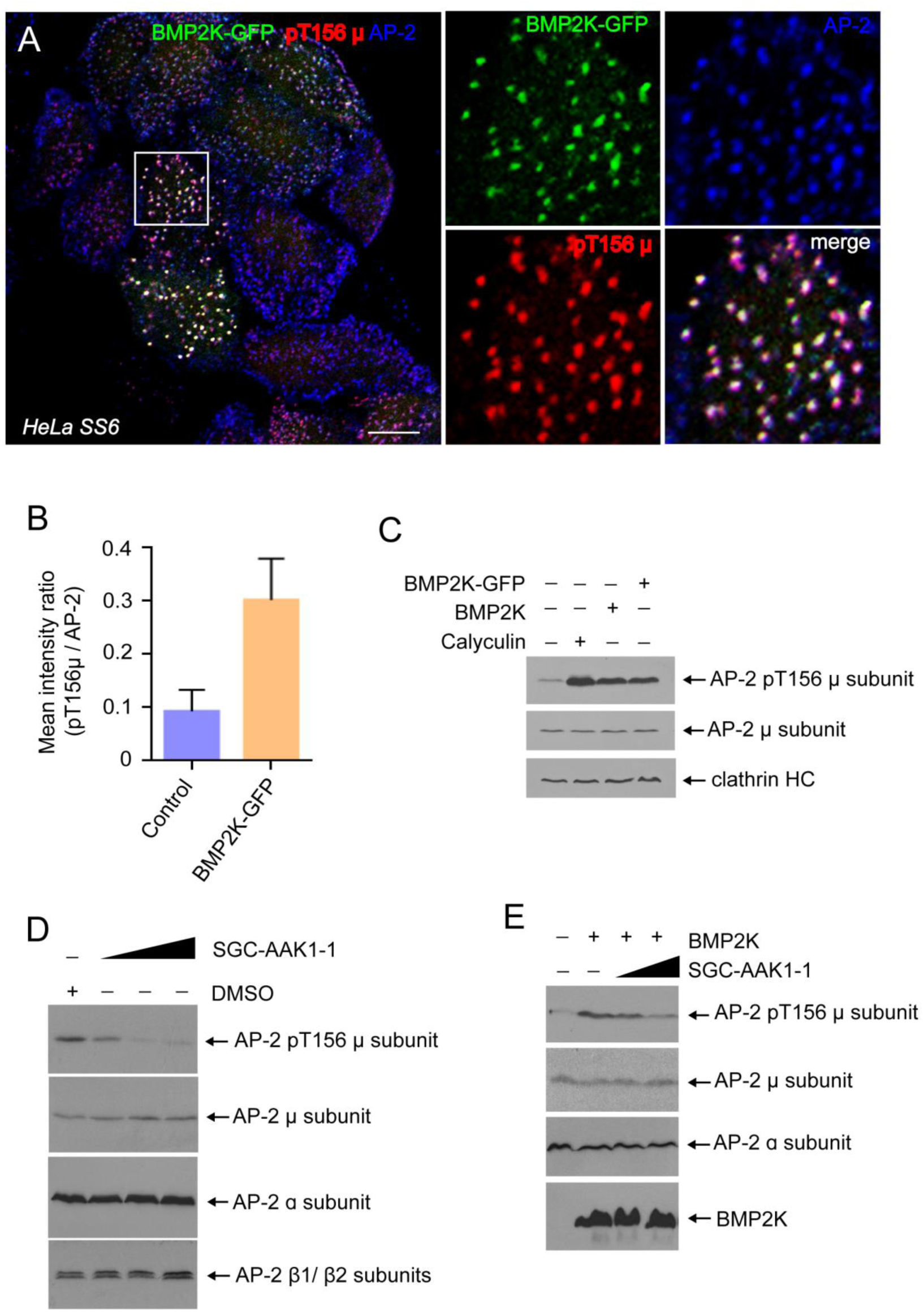
BMP2K is a *bona fide* AP-2 kinase in cells. (**A**) Representative confocal optical section of the basal plane of HeLa SS6 cells transfected with BMP2K-GFP (green), fixed and immunolabeled for phosphorylated AP-2 (pT156µ, red) and AP-2 (blue). Colour separated channels of the boxed region are shown on the right. Scale bar-10 µm. (**B**) Bar graphs comparing the ratio of mean fluorescence intensities of phosphorylated AP-2 (pT156µ) to total AP-2 in CCPs in transfected (BMP2K-GFP) versus untransfected (Control) HeLa SS6 cells. Data represent mean SD of 29 cells analysed from 3 independent experiments. **(C)** Immunoblot analysis of whole cell lysates from HeLa SS6 either DMSO or calyculin treated or transfected with BMP2K or BMP2K-GFP as indicated. Blots are probed with the indicated antibodies. **(D)** Immunoblot analysis of whole cell lysates from HeLa SS6 treated with either DMSO or increasing concentrations of SGC-AAK1-1 for 2 hours. Blots are probed with the indicated antibodies. **(E)** Immunoblot analysis of whole cell lysates from HeLa SS6 either mock transfected or transfected with BMP2K-GFP and treated with either DMSO or increasing concentrations of SGC-AAK1-1 for 2 hours. Blots are probed with the indicated antibodies.

### BMP2K localizes to CCPs through extended C-terminus

BMP2K positive CCPs also contain FCHO binding partners, clathrin-associated SNARE adaptor, HIV-1 Rev-binding protein (HRB) (Pryor et al., 2008) (**Figure 3- figure supplement 1A**) and pioneer adaptor complex protein, intersectin (Henne et al., 2010; Mayers et al., 2013) (**Figure 3- figure supplement 1B**), further confirming endocytic association of kinase. Yet, distribution of BMP2K-GFP appears slightly different from that of GFP-FCHO1. In abundantly expressing cells, larger structures that concentrate more AP-2 (**Figure 3A**) are seen in both, but membrane-tethered population evident with higher expression levels of GFP-FCHO1 (**Figure 3B**) (Umasankar et al., 2012) is not observed with higher amounts of BMP2K-GFP. This likely indicates differential mode of CCP recruitment and stabilization for FCHO and BMP2K. We used deletion mutants to delineate mode of CCP recruitment of BMP2K. BMP2K follows domain architecture typical of the parental NAK-consisting of NH_2_-terminal kinase domain (KD) and an extended COOH-terminus with proximal glutamine/histidine (Q/H) rich region and distal intrinsically disordered region (CT) (**Figure 3C**). Deletion of CT (BMP2KΔCT) completely abolishes CCP localization (**Figure 3D**). Yet, deletion of KD and Q/H together, retaining CT alone (BMP2KCT) or deletion of Q/H alone retaining KD and CT (BMP2KΔQ/H) shows reduced localization at CCPs (**Figure 3E** and **F**). Strikingly, deletion of KD alone (BMP2KΔKD) completely restores CCP localization (**Figure 3G**), suggesting localization happens through co-ordination of Q/H and CT regions independent of KD. This is in contrast to the structured, domain-dependent CCP targeting of FCHO (Umasankar et al., 2012).

**Figure 3:**
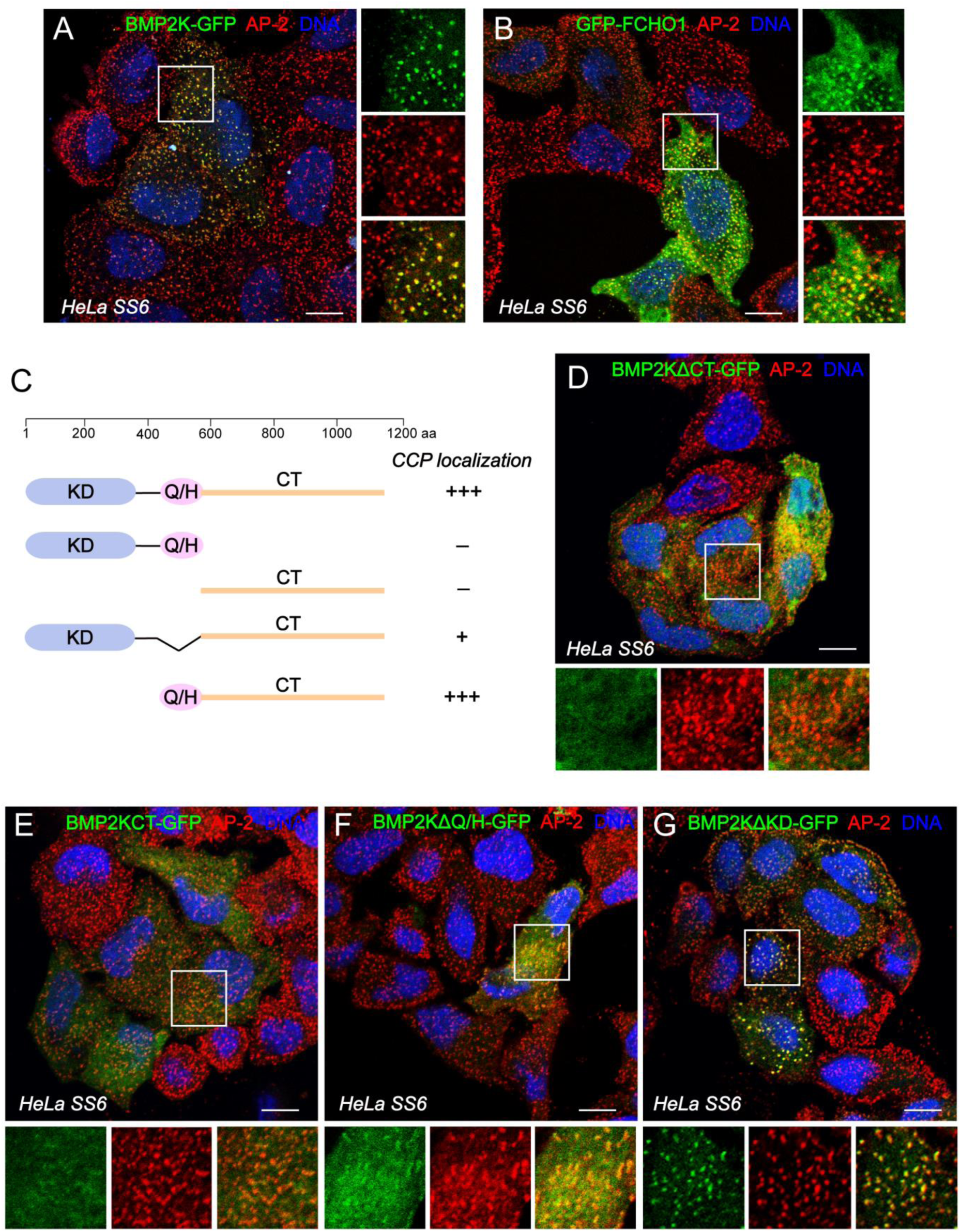
CCP localization of BMP2K *via* C-terminal region. (**A and B**) Representative confocal optical sections of the basal plane of HeLa SS6 cells transfected with BMP2K-GFP (green) (**A**) or GFP-FCHO1 (green) (**B**), fixed and immunolabeled for AP-2 (red). Colour separated views of the boxed regions are shown on the right. (**C**) Schematic depiction of the overall domain organization of BMP2K (1-1161) and relative locations of the various C-terminal GFP tagged truncation and deletion mutants of BMP2K tested. Efficiency of CCP localization of each mutant is marked. (**D-G**) Representative confocal images of HeLa SS6 cells transiently transfected with BMP2KΔCT-GFP (**D**), BMP2KCT-GFP (**E**), BMP2KΔQ/H-GFP (**F**) and BMP2KΔKD-GFP (**G**). Fixed cells were stained for AP-2 and DNA. Colour separated channels of boxed regions are shown under each image. Scale bar-10 µm.

### BMP2K binds to AP-2 and clathrin

To identify endocytic interactions involved in BMP2K recruitment to CCP, we performed GST-pull-down assays with HeLa SS6 cell extracts. CT contains known and predicted short linear motifs (SLiMs) distributed across proximal (CT#1) and distal (CT#2) segments that can potentially engage AP-2 appendages, clathrin and muniscins (**Figure 4A**). Accordingly, both GST-CT#1 and GST-CT#2 affinity purify α and β subunits of AP-2 from cell lysate (**Figure 4B**). EPS15 C-terminus (595-896) binds directly to appendages through multiple low affinity DPF motifs and indirectly to clathrin through AP-2 (**Figure 4B**, Benmerah et al., 1998). Compared to EPS15, CT shows weaker binding to AP-2 and to clathrin in our pull-down experiments despite having an array of potential binding motifs. Additionally, EPS15 contains three tandem, optimally spaced DPFXXDPF consensus motifs that engage FCHO for CCP initiation (Ma et al., 2016). We identify nine, similar DXFXXXPF consensus motifs scattered throughout BMP2K-CT (**Figure 4A**); one of these motifs can directly engage FCHO2 (Mulkearns et al., 2012). Yet, BMP2K does not show any obvious interaction with FCHO (**Figure 4B**). This confirms the importance of spatial arrangement of SLiMs or redundant mechanisms in mediating diverse interactions to build clathrin-coat (Ma et al., 2016). Surprisingly, Q/H region, despite showing definitive role in CCP targeting, shows negligible AP-2 or clathrin binding and is comparable to KD (**Figure 4C**).

**Figure 4:**
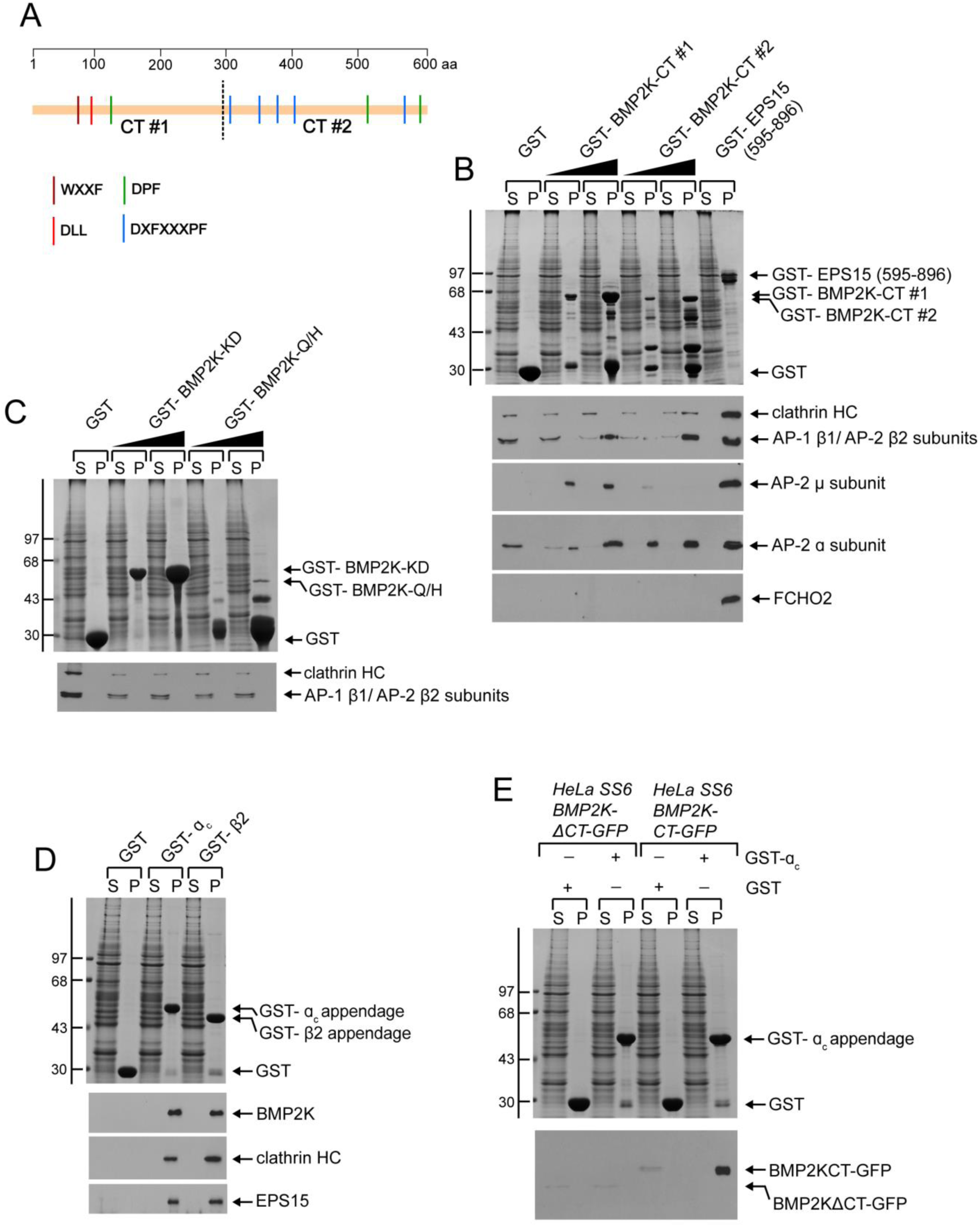
Reciprocal interactions of BMP2K and AP-2. (**A**) Cartoon diagram showing distribution of various short linear interacting motifs (SLiMs) on the distal (CT#1) and proximal (CT#2) segments of the BMP2K C-terminus. (**B-D**) Biochemical pull-down assays of Triton-X-100 lysates of HeLa SS6 cells with fixed concentrations of GST (**B and C**), GST-EPS15 (595-896) (**B**) increasing concentrations of GST-BMP2K-CT #1, GST-BMP2K-CT #2 (**B**), GST-BMP2K-KD, GST-BMP2K-Q/H (**C**) or with equal concentrations of GST, GST-α_c_ and GST-β2 appendages (**D**). (**E**) Biochemical pull-down assays with equal concentrations of GST and GST-α_c_ appendage and equal amounts of Triton-X-100 lysates of HeLa SS6 cells expressing either BMP2KCT-GFP or BMP2KΔCT-GFP. After pull down assays, samples of supernatant (S) and pellet (P) fractions were analysed on SDS-PAGE and either stained with Coomassie blue along with molecular mass standards (in kDa) on the left as indicated (**B-E**, top panels) or transferred to nitrocellulose membrane and immunoblotted for the indicated proteins using specific antibodies (**B-E**, bottom panels).

### AP-2-dependent BMP2K stabilization at CCPs is pivotal for phosphorylation

AP-2 interacts with accessory endocytic proteins through α and β appendages to regulate their recruitment to CCP (Mishra et al., 2004; Praefcke et al., 2004; Keyel et al., 2008). Accordingly, in reciprocal pull-down assays, both GST-α and – β appendages affinity purify endogenous BMP2K from HeLa lysate (**Figure 4D**). In addition, appendage engages CT alone and fails to bind ectopically expressed BMP2KΔCT, further underscoring the importance of CT-AP-2 interaction (**Figure 4E**).

We checked if functional AP-2 is indeed required for BMP2K stabilization at CCP. siRNA targeting AP-2 α subunit efficiently suppresses AP-2 function in HeLa cells (Motley et al., 2003, Hinrichsen et al., 2003). Immunofluorescence with AP-2 monoclonal antibody AP.6 clearly distinguishes AP-2 depleted cells from adjacent normal cells in the same field (**Figure 5A**). In addition, fluorescent transferrin (endocytosis assay-protocol #1) completely stalls at the cell surface of AP-2 depleted cells and fails to internalize into peripheral endosomes confirming functional inactivation of AP-2 in these cells (**Figure 5B**). Interestingly, in AP-2 depleted cells (**Figure 5C**), BMP2K-GFP fails to localize to CCPs and appears diffused throughout cytosol (**Figure 5D**). Immunoblots reveal knockdown of α subunit abolishes AP-2 complex formation and thus causes degradation of β, µ and σ sub-units. Interestingly, steady state levels of endogenous BMP2K also sharply decline in AP-2 depleted cells indicating destabilization and degradation of kinase (**Figure 5E**). Further, we checked if AP-2 phosphorylation is regulated through CCP localization of the kinase. As expected, steady-state intensity of pT156 µ considerably increases in CCPs populated with BMP2K-GFP (**Figure 5F**). However, phosphorylation status of AP-2 remains unaffected in BMP2KΔCT-GFP (that cannot engage AP-2) expressing cells, despite having functional KD. (**Figure 5G**). Together, it implies that membrane AP-2 stabilizes kinase at growing CCPs for catalysis.

**Figure 5:**
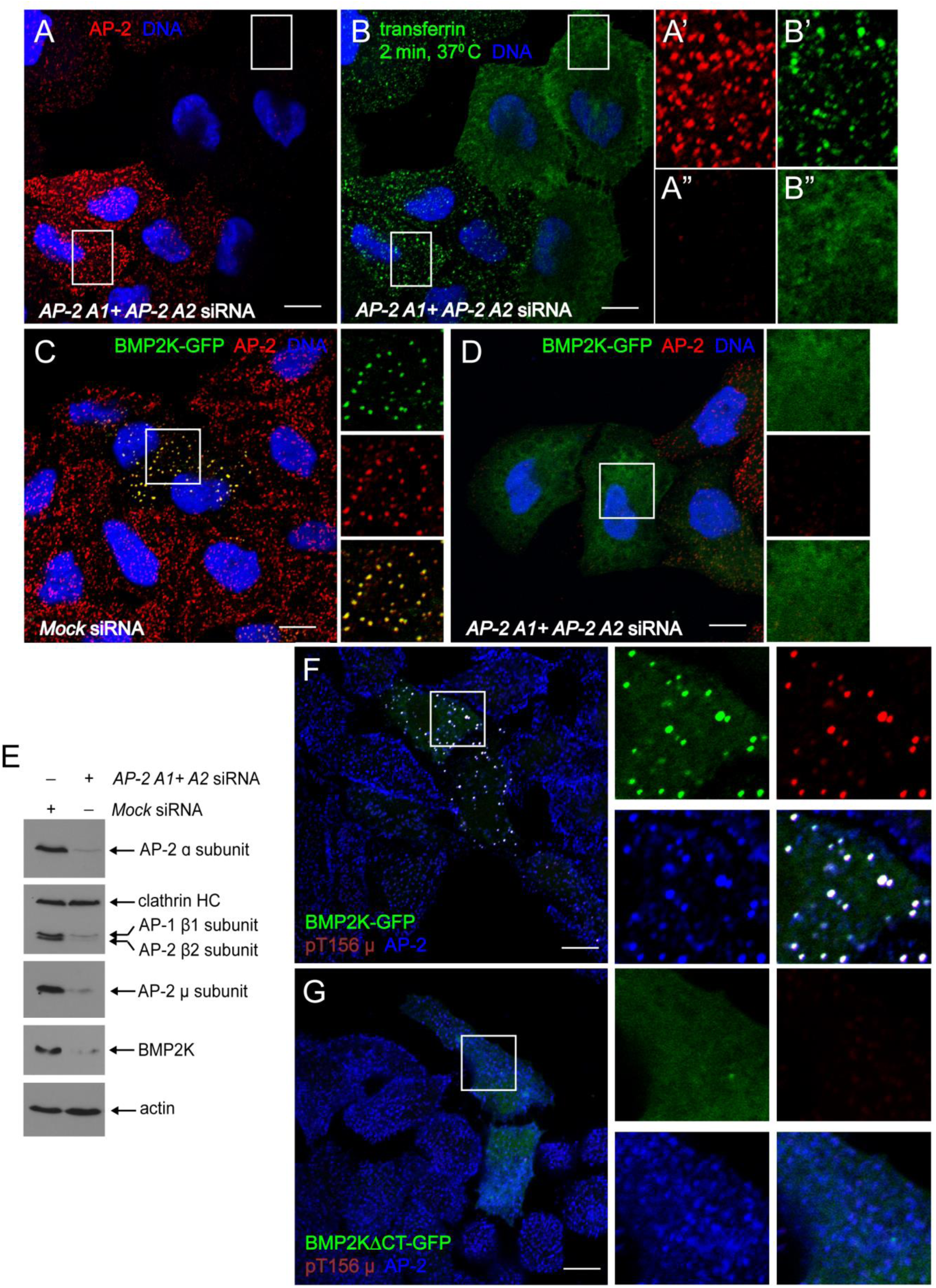
AP-2 interaction is important for CCP recruitment, stabilization and function of BMP2K. (**A-B”**) Representative confocal optical sections of HeLa SS6 cells transfected with siRNA targeting AP-2 α subunit (A1 and A2). Cells were stained with anti-AP-2 α subunit mAb AP.6 (red) as indicated (**A**), after 2 minute pulse with Alexafluor488-transferrin (green) (**B**). Boxed regions of AP-2 containing (**A’, B’**) and AP-2 depleted (**A”, B”**) cells in the same field (**A, B**) are shown as enlarged and colour separated images on the right. (**C** and **D**) Representative confocal images of HeLa SS6 cells expressing BMP2K-GFP. Mock transfected (C) or AP-2 α siRNA transfected (D) cells were labeled for AP-2 and DNA as indicated (**C, D**). Zoomed, colour separated and merged images of the boxed regions are displayed on the right. (**E**) Comparative immunoblot analysis of whole cell lysates of either mock transfected or AP-2 α siRNA transfected HeLa SS6 showing extent of AP-2 and BMP2K depletion. Blots are probed with the indicated antibodies. (**F** and **G**) Representative confocal sections of HeLa SS6 cells transiently transfected with either BMP2K-GFP (**F**) or BMP2KΔCT-GFP (**G**) (green) and immunolabeled for phosphorylated AP-2 (pT156µ, red) and AP-2 (blue). Colour separated channels of the boxed region are enlarged and shown on the right. Scale bar-10 µm.

### BMP2K regulates CCP morphology and cargo uptake

Silencing BMP2K using specific pool of siRNAs did not cause any discernible alterations in CCP morphology or cargo uptake in our experiments. This is consistent with lack of CME phenotype in AAK1 knockdown cells in earlier studies suggesting functional redundancies in AP-2 kinases (Conner and Schmid, 2003). Hence, we took a pharmacological approach to functionally inactivate BMP2K/AAK1 in HeLa SS6 cells. Our biochemical experiments reveal that 10 µM SGC-AAK1-1 (NAK inhibitor) shows maximum inhibition of AP-2 phosphorylation in 2 hours beyond which it remains unchanged (**Figure 2D**), hence 10 µM was selected for inhibition experiments. BMP2K inhibited parental HeLa SS6 cells (*BMP2K i*) show visible alterations in the steady-state CCP morphology. Quantitative analyses of CCP density and distribution in mock cells show predominantly small, spherical structures (**Figure 6A and Figure 6- figure supplement 1A**) ranging 0.2-0.6 µm^2^ size that occupy ∼ 40% of total membrane bound AP-2 (**Figure 6D and E**). *BMP2K i* cells exhibit a clear shift in the distribution and density of CCPs toward enlarged, sparsely distributed, brighter structures (**Figure 6B and Figure 6- figure supplement 1B**), above 0.6 µm^2^ size that occupy ∼ 60% of total AP-2 (**Figure 6D and E**) on the membrane. These spatially remodelled AP-2 positive structures in BMP2K *i* cells morphologically resemble flat, planar, clathrin-coated plaques in 1.E cells (**Figure 6C**, Umasankar et al., 2014). To see if altered CCP morphology in *BMP2K i* cells translates into functional defects similar to 1.E cells, we compared kinetics of transferrin internalization in mock, *BMP2K i* and 1.E cells using endocytosis assay protocol #1. A flux of fluorescent transferrin in mock cells separates AP-2 positive CCPs from peripheral early endosomes in 2 minutes (**Figure 6A’**) and from large juxtanuclear recycling endosomes in 10 minutes (**Figure 6- figure supplement 1A-A”**). Consistent with our earlier reports, the bulk of transferrin appears diffused throughout the cell surface with very low amounts in peripheral endosomes suggesting cargo sorting defects in 1.E cells (**Figure 6C’ and C”**). Strikingly, in *BMP2Ki* cells, bulk of transferrin co-localizes with AP-2 positive structures (**Figure 6B”)** with reduced internalization into endosomes in 2 minutes (**Figure 6B’**). In addition, after 10 minutes of pulse, the fluorescent transferrin puncta appears more distributed in peripheral early endosome compartments instead of juxtanuclear recycling endosome compartments in *BMP2K i* cells (**Figure 6- figure supplement 1B’** and **B”**). Collectively, these findings suggest persistent CME delay in the absence of functional kinase. However, transferrin diffusion at cell surface seen in 1.E cells is not clearly evident in *BMP2K i* cells likely indicates temporally distinct functions for FCHO and BMP2K in CME or, in part, reports differences in acute (pharmacological) or chronic (genome-editing) interference defects in CME. Together, these findings argue a late endocytic function for kinase, which is consistent with recent reports indicating role of AP-2 phosphorylation in CCP maturation (Wrobel et al., 2019).

**Figure 6:**
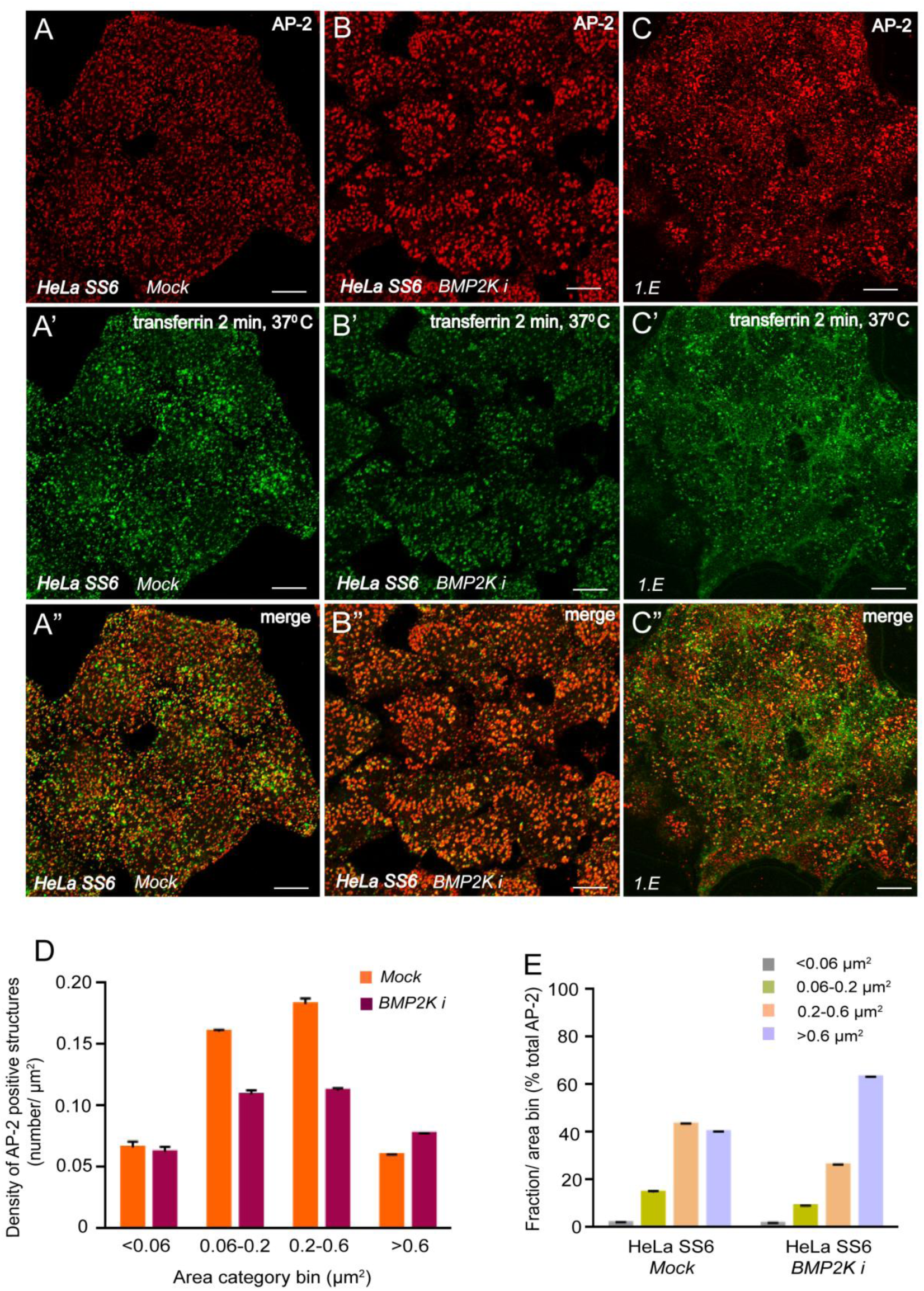
Functional inactivation of BMP2K and FCHO shows temporally distinct CME phenotypes in HeLa cells. (**A-C”**) Colour separated and merged channels from representative confocal images of HeLa SS6 cells treated with 10 µM DMSO (*Mock*) (**A-A”**), 10 µM SGC-AAK1-1 inhibitor (*BMP2K i*) (**B-B”**) for 2 hours or 1.E cells (**C-C”**), pulsed with Alexafluor488-transferrin (green), immunostained for AP-2 (red) to compare CCP morphology and endocytic efficiency. Scale bar-10 µm. (**D and E**) Bar graphs represent quantification of number of AP-2 positive structures in each area category bin (previously classified based on size and dynamics as explained in Ma et al., 2016) per µm^2^ cellular area (D) or percentage of total AP-2 occupied by each bin size (color-coded) (E) in *Mock* or inhibitor treated (*BMP2K i*) HeLa SS6 cells. Data represent mean SD from total cellular areas of mock (30589.55 µm^2^) and *BMP2K i* (28527.73 µm^2^).

**Figure 7:**
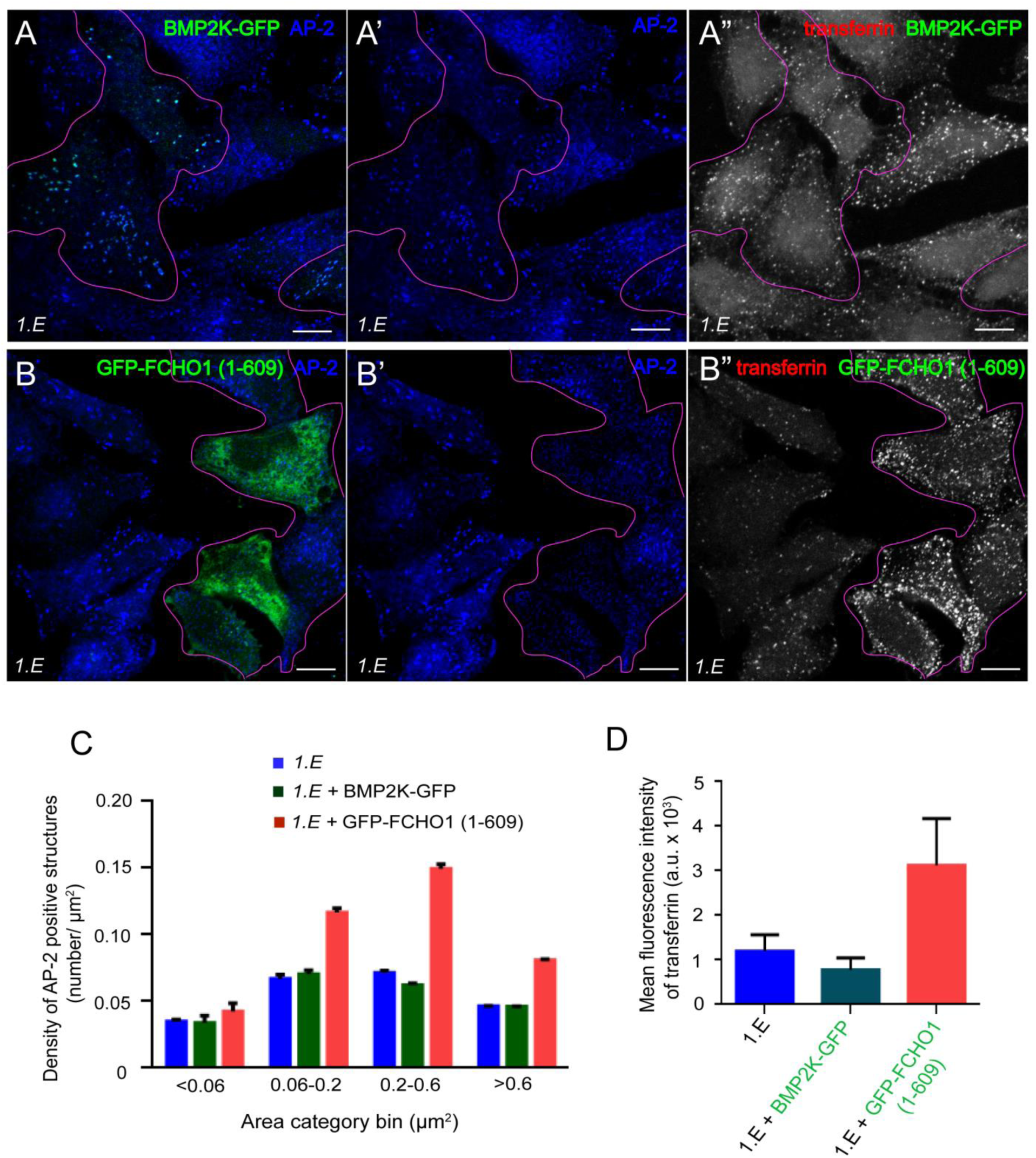
Differential effects of BMP2K and FCHO linker expression on CCP morphology and function in 1.E cells. (**A-B”**) Representative confocal images of 1.E cells transiently transfected with BMP2K-GFP (**A-A”**) or GFP-FCHO1 (1-609) (**B-B”**), pulsed with Alexafluor568-transferrin (pseudocolored grey) for 2 minutes, surface stripped and allowed to internalize for another 2 minutes (protocol#3), and immunostained for AP-2 (blue). Single basal optical planes (**A, A’, B, B’**) or merged stack of several optical planes along Z-axis (**A”, B”**) are displayed. Scale bar-10 µm. BMP2K-GFP and GFP-FCHO1 (1-609) expressing cells are outlined (purple) in corresponding optical sections. (**C and D**) Quantification of number of AP-2 positive structures in each area category bin per µm^2^ cellular area (C) or mean fluorescence intensity of internalized transferrin (D) in 1.E, 1.E + BMP2K-GFP and 1.E + GFP-FCHO1 (1-609). Graphs show mean SD from total cellular areas analysed for 1.E (12874.71 µm^2^), 1.E + BMP2K-GFP (6740.73 µm^2^) and 1.E + GFP-FCHO1 (1-609) (3716.3 µm^2^).

### FCHO-BMP2K feed-forward axis in HeLa cells

We surmise if BMP2K drives AP-2 functions in the absence of FCHO, then re-expression of BMP2K in 1.E cells could restore normal CCP morphology and function. Ectopic expression of BMP2K-GFP in 1.E cells marks AP-2 positive structures on membrane (**Figure 7A**) but does not alter CCP morphology or distribution; large bright plaques still persist in these cells (**Figure 7A’ and C**). To check if BMP2K positive structures in 1.E cells are functional, we assessed cargo uptake using endocytosis assay protocol #3 that clearly discerns small changes in internalization kinetics (Reis et al., 2015; Ma et al., 2016). Negligible difference is seen in the amount of internalized transferrin in BMP2K-GFP transfected cells compared to surrounding untransfected 1.E cells; suggesting BMP2K activation fails to rescue AP-2 functions in the absence of FCHO (**Figure 7A”, D**). However, complete correction of CCP morphology and cargo uptake defects occurs with expression of a membrane-binding segment of FCHO1 GFP-FCHO1 (1-609) containing the linker region (316-467) that interacts with AP-2 core (Umasankar et al., 2014; Hollopeter et al., 2014). Within linker transfected 1.E cells, CCP population density shifts predominantly toward 0.2-0.6 µm^2^ sized AP-2 positive structures (**Figure 7B**, **B’ and C**). Linker transfected cells also show considerable increase in the amount of intracellular transferrin further confirming rescue of CME defects through allosteric activation of AP-2 (**Figure 7B” and D**).

We asked if FCHO could restore normal AP-2 phosphorylation in 1.E cells by activating functional kinase. Similarly to linker, reexpression of GFP-FCHO1 in 1.E cells restores normal AP-2 distribution on the membrane (**Figure 8A, A’** and **C-F**). Interestingly, mean pT156 µ intensity increases ∼ 2.5 fold in AP-2 positive structures containing GFP-FCHO1 as compared to the surrounding AP-2 positive plaques devoid of GFP-FCHO1 in the same field of view (**Figure 8B, B”, G-K**). Together, these findings suggest AP-2 functions through FCHO-BMP2K feed-forward axis in mammalian cells.

**Figure 8:**
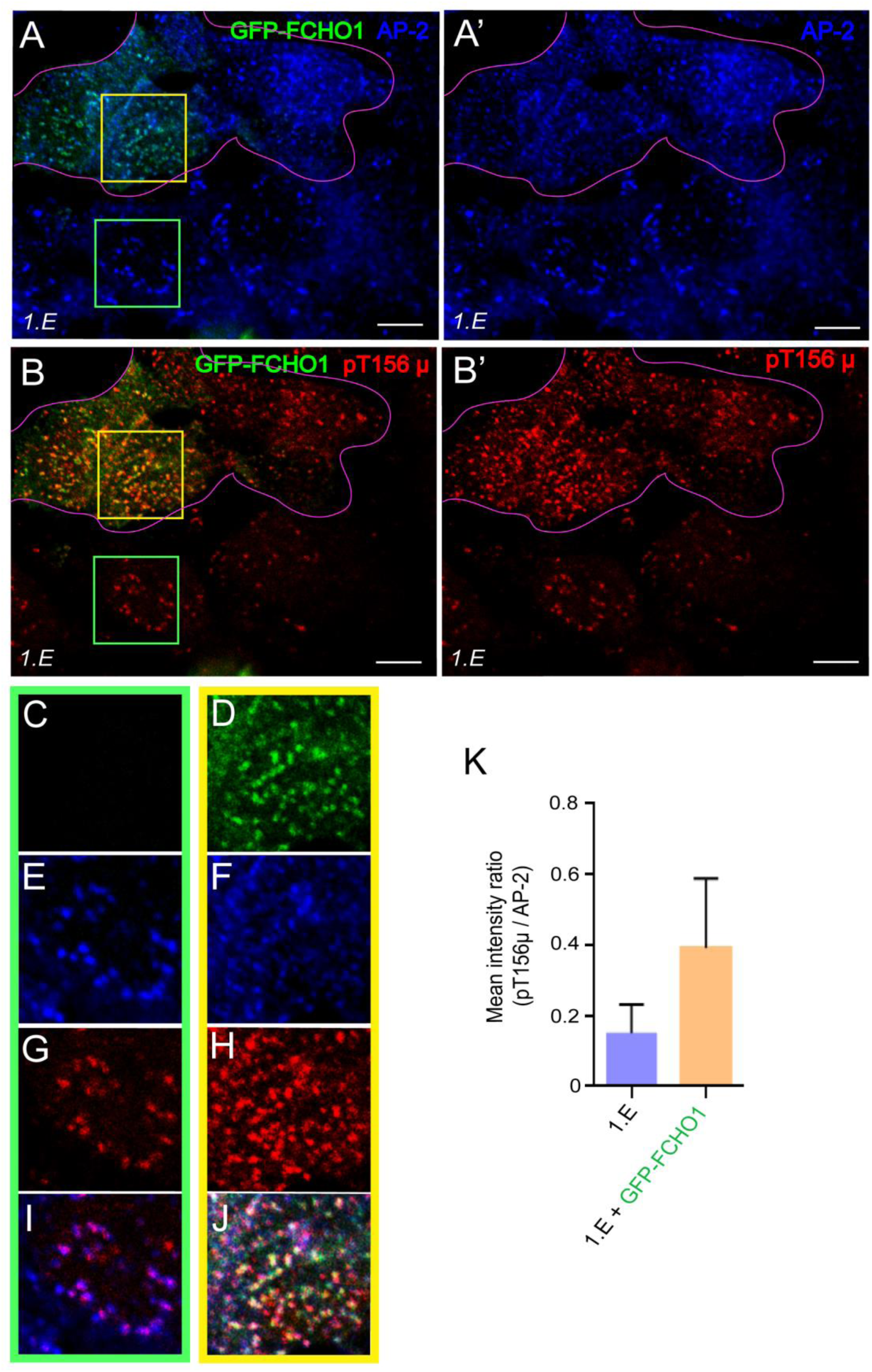
FCHO expression restores AP-2 phosphorylation in 1.E cells. (**A-J**) Representative single confocal optical sections of 1.E cells transiently transfected with GFP-FCHO1, fixed and immunolabeled for AP-2 (**A and A’**) and phosphorylated AP-2 (pT156µ) (**B and B’**). Scale bar-10 µm. GFP-FCHO1 expressing cells are outlined (purple) in corresponding optical sections. Colour separated and merged channels of the boxed regions from transfected and untransfected areas are displayed (**C-J**). (**K**) Bar graphs showing increase in mean fluorescence intensity ratio of phosphorylated AP-2 (pT156µ) to total AP-2 in CCPs in GFP-FCHO1 transfected 1.E cells compared to untransfected 1.E cells. Data represent mean SD from 14 cells.

### FCHO misexpression phenocopies BMP2K gain-of-function *in vivo*

Next, we investigated if FCHO-BMP2K axis functions at organismal level for AP-2 activation, by using zebrafish (*Danio rerio*) embryos. Single genes each code for *Dr* FCHO1 (*fcho1*), *Dr* FCHO2 (*fcho2*) and *Dr* AP-2 α (*ap2a1*) subunit (Umasankar *et al*, 2012). However, *Dr* BMP2K and *Dr* AAK1 are encoded by two genes each that might have formed due to teleost-specific genome duplication during evolution (Howe et al., 2013). Whole mount *in situ* hybridization analyses using riboprobes designed from common kinase domain of *bmp2k1* and *bmp2k2* reveal maternal deposition and ubiquitous expression of kinase in all developmental stages till 24 hour post-fertilization (hpf) (**Figure 9- figure supplement 1A**). Spatio-temporal expression pattern of BMP2K is identical to that of *ap2a1* (**Figure 9- figure supplement 1B**) and *fcho1/2* transcripts; (Umasankar *et al*, 2012) and is in stark contrast to that of *dusp6* (dual specificity phosphatase-6), a non-endocytic transcript that is not maternally deposited but shows specific domains of expression during zygotic phase of development (**Figure 9- figure supplement 1C**) (Tsang et al., 2004; Molina et al., 2009). This spatio-temporal expression pattern shows CME is functional in all stages of embryonic development.

**Figure 9:**
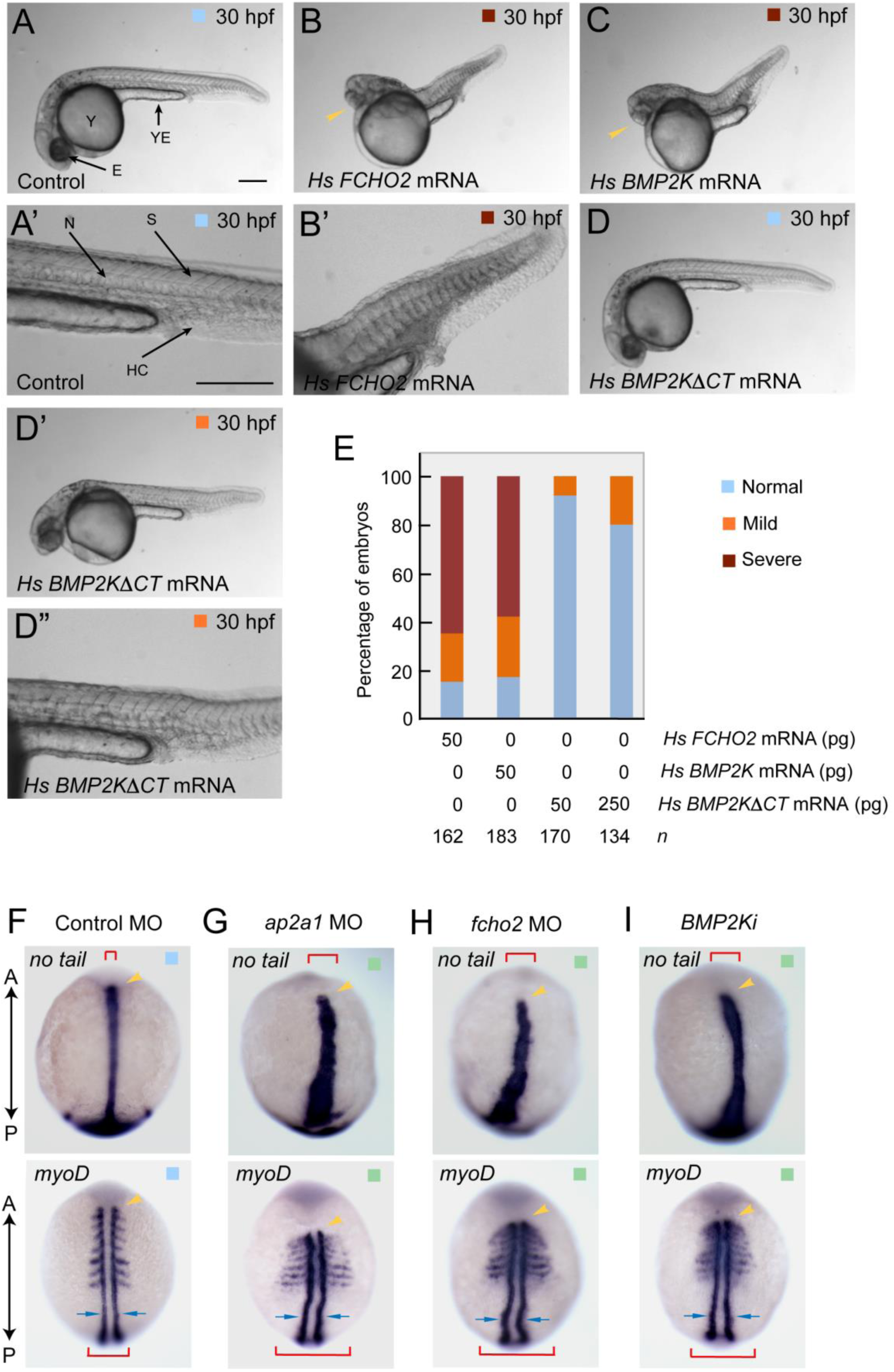
FCHO, AP-2 and BMP2K co-ordinate CME functions during embryonic development. (**A-D”**) Representative bright field live images of control (**A**), 50 pg human (*Hs*) *FCHO2* mRNA (**B**), 50 pg *Hs BMP2K* mRNA (**C**), 50 pg *Hs BMP2KΔCT* mRNA (**D**) and 250 pg *Hs BMP2KΔCT* mRNA (**D’**) injected *D. rerio* embryos at 30 hours post-fertilization (hpf). Enlarged images focusing the caudal region of embryos (**A’, B’ and D”**). Black arrows indicate different structures in the embryo E: eye, Y: yolk, YE: yolk extension, N: notochord, S: somites, HC: haematopoetic cells. Cyclopic eye phenotype is denoted with yellow arrow heads in **B** and **C**. Colour coded as in **E**. Scale bar-250 µm (**E**) Bar graphs showing quantification of normal (pale blue), mild (orange) and severe (brown) mRNA overexpression phenotypes. *n*: number of embryos analysed. (**F-I**) Representative whole-mount *in situ* images showing mRNA expression patterns of *no tail* (top panels) and *myoD* (bottom panels) in 6-somite stage *D.rerio* embryos injected with control (**F**), *ap2* morpholino targeting α subunit (*ap2a1 MO*) (**G**), *fcho2* morpholino (*fcho2 MO*) (**H**) or treated with SGC-AAK1-1 (*BMP2Ki*) (**I**) as indicated. A: anterior, P: posterior. Red brackets and yellow arrow heads indicate lateral and axial expression territory of markers respectively. Blue arrows denote adaxial cell boundary in *myoD* stained embryos. Colour coded as in figure supplement-**2C**.

Ectopic expression of *Hs FCHO2* mRNA in *D. rerio* embryos causes severe developmental defects. At 30 hpf, control embryos show normal axial patterning and body plan (**Figure 9A** and **A’**). But, 65 % of *Hs FCHO2* mRNA injected embryos exhibit cyclopia (fusion of two eyes due to lack of separating floor plate), shortened and upward twisted A-P axis with reduced yolk extension, disrupted notochord tissue, misshapen somites and accumulated caudal hematopoietic cells (classified as severe) (**Figure 9B, B’** and **E**). 20% of the injected embryos show only mild notochord and somite defects without cyclopia or upward twisted body axis (classified as mild) and 15% appear similar to control (classified as normal) (**Figure 9E**). These phenotypes closely resemble *Hs FCHO1* mRNA misexpression phenotypes noticed in our earlier *in vivo* studies (Umasankar *et al*, 2012). Strikingly, ectopic expression of *Hs BMP2K* mRNA phenocopies FCHO misexpression with 58% severe, 25% mild and 17% normal embryos (**Figure 9C** and **E**). However, expression of BMP2K mutant that cannot engage AP-2 (*Hs BMP2KΔCT*), fails to elicit severe embryonic defects (**Figure 9D** and **E**). Only 20% of embryos injected with 5 times higher concentration of *Hs BMP2KΔCT* mRNA exhibit mild phenotypes further confirming *in vivo* significance of BMP2K-AP-2 interaction (**Figure 9D’, D”** and **E**). This suggests that hyperactivation of AP-2 by misexpressing FCHO or BMP2K could increase CME rates leading to multiple but similar defects during embryogenesis. Thus, gain-of-function studies confirm FCHO and BMP2K are functionally linked in vertebrates.

### Functional FCHO-BMP2K axis *in vivo*

Next, we compared loss-of-function phenotypes of FCHO and BMP2K with respect to specific AP-2 functions during development. Since AP-2 knockout is embryonically lethal in vertebrates (Mitsunari et al., 2005), we used anti-sense morpholino approach to transiently knockdown *ap2a1* in *D. rerio* embryos. We have used this approach previously to functionally inactivate AP-2 and other endocytic proteins in developing embryos. In addition, target specificity and knockdown efficiency of these morpholinos have been validated previously by control/ mismatch morpholinos, GFP-tagged target silencing and functional rescue experiments (Umasankar et al., 2012).Translational silencing of *ap2a1* causes severe defects in morphogenetic cell movements during embryonic gastrulation. Through an unbiased morphological screen, we classified these gastrulation phenotypes into two categories. Category-1 embryos are severely affected and show slow and incomplete epiboly during early gastrulation. Compared to normally developing control morpholino-injected embryos (**Figure 9- figure supplement 2A** and **A’**), category-1 morphants perish during blastopore closure at ∼ 9-10 hpf due to dysmorphic extrusion of yolk (**Figure 9- figure supplement 2B** and **B’**). Embryonic culture of blastomeres from category-1 embryos reveals defects in cell adhesion. Unlike control cells that exhibit typical fibroblast-like appearance (**Figure 9- figure supplement 2D**), majority of *ap2a1* morphant cells appear rounded and dissociated, indicative of failure to adhere and spread out on laminin containing extracellular matrix (**Figure 9- figure supplement 2E**). Endocytosis promotes epiboly by removing the yolk-cell membrane just ahead of the advancing blastoderm (Solnica-Krezel., 2006) and also by recycling cell adhesion protein, E-cadherin which contains di-leucine sorting signal for AP-2 (Song et al., 2013). Consistent with this idea, *ap2a1* category-1 embryos phenocopy dynamin mutants (Lepage et al., 2014) Rab5 morphants (Kenyon et al., 2015) and *half baked*/E-cadherin mutants (Kane et al., 2005).

Category-2 embryos undergo normal epiboly but display convergent-extension (CE) defects during late gastrulation. WISH analyses using molecular markers *no tail* (for notochord cells) and *myoD* (for somite and adaxial cells) reveal chordamesoderm cells undergoing normal CE in all control morpholino injected embryos (**Figure 9F** and **Figure 9- figure supplement 2C**). In vertebrates, CE is governed by non-canonical Wnt-PCP signaling that co-ordinate cell movements *via* Rho GTPase-actin axis (Tada and Heisenberg, 2012; Roszko et al., 2009). Strength and duration of Wnt-PCP signaling rely on endocytosis of Wnt-signalosome complex consisting of Wnt (ligand)-Frizzled (receptor)-Dishevelled (adaptor containing tyrosine sorting signal) through functional AP-2 (Yu et al., 2007; Ohkawara et al., 2011). Accordingly, *ap2a1* morphants display severe CE defects; territory of both *no tail* and *myoD* expressing cells shorten across A-P axis and expands laterally, reminiscent of Wnt-PCP mutants (Hammerschmidt et al., 1996; Hagemann et al., 2014) (**Figure 9G**). Surprisingly, embryos injected with *fcho2* morpholinos develop normally with no obvious epiboly defects (category-1). However, 79% *fcho2* morphants phenotypically mimic *ap2a1* category-2 embryos showing CE defects with respect to *no tail* and *myoD* expression pattern (**Figure 9H** and **Figure 9- figure supplement 2C**). These findings suggest redundant mechanisms for AP-2 activation during embryogenesis (Umasankar et al., 2012). Pharmacological approaches suppress BMP2K function in developing embryos. Consequently, gastrulating embryos treated with SGC-AAK1-1 (*BMP2Ki*) show clear CE defects (60% embryos fall into category-2) analogous to *fcho2* and *ap2a1* morphants (**Figure 9I** and **Figure 9- figure supplement 2C**). Together, these findings certify FCHO-BMP2K axis functions across vertebrates to regulate AP-2.

## Discussion

GTPase-independent initiation mechanisms impose multiple layers of regulation in CME and mark it different from other vesicular transport pathways (Ren et al., 2013; Nie et al., 2003; Umasankar et al., 2014; Hollopeter et al. 2014; Godlee and Kaksonen, 2013). Activation of AP-2 through hierarchical, allosteric conformational modifications has been proposed a major regulatory mechanism that drives CCV biogenesis (Kadlecova et al., 2017; Mettlen et al., 2018). Additionally, interplay of various checkpoints at multiple stages of CCV biogenesis-initiation, cargo selection, maturation, and fission are also suggested to ensure quality control of endocytosis such as incorporation of cargo into CCP before maturation, disassembly of defective coated-pits *etc*. (Loerke et al., 2009; Mettlen et al., 2009; Aguet et al., 2013). Recent investigations revealing functionally distinct conformations of AP-2 *viz*; closed (AP-2_close_), open (AP-2_open_), open+ (AP-2_open+_) and phosphorylated open+ (AP-2_phospho_) provide valuable clues about spatial subunit rearrangements in AP-2 activation. (Collins et al., 2002; Kelly et al., 2008; Jackson et al., 2010; Kelly et al., 2014; Wrobel et al., 2019).

Our study proposes a model that consolidates recent findings and provides novel *in vivo* mechanisms to define AP-2 activation (**Figure 10**). Here, we have identified two allosteric AP-2 activators *viz*; FCHO and BMP2K that entropically alter conformational equilibrium (Buchenberg et al., 2017; Wieteska et al., 2018) from cytosolic inactive AP-2 to membrane-bound phosphorylated AP-2 by co-ordinating a feed-forward axis. We also show that this allosteric axis enables vectorial progression of CCV biogenesis. Spatial proteome analyses denote ∼80 % of AP-2 is membrane bound in mammalian cells (Itzhak et al., 2016). Yet, a large pool of AP-2 remains unphosphorylated in FCHO knockout cells. These findings argue that FCHO must be functioning together with EPS15/R to allosterically drive the equilibrium from cytosolic inactive AP-2_close_ to membrane active AP-2_open_ / AP-2_open+_ (Ma et al., 2016; Traub, 2019). This equilibrium is further driven toward CCP maturation by downstream kinase that phosphorylates open+ state of AP-2 (Wrobel et al., 2019). Thus, absence of FCHO or kinase would perturb this axis and reverse the equilibrium leading to defects in cargo sorting, clathrin assembly and maturation (this study and Umasankar et al., 2014).

**Figure 10:**
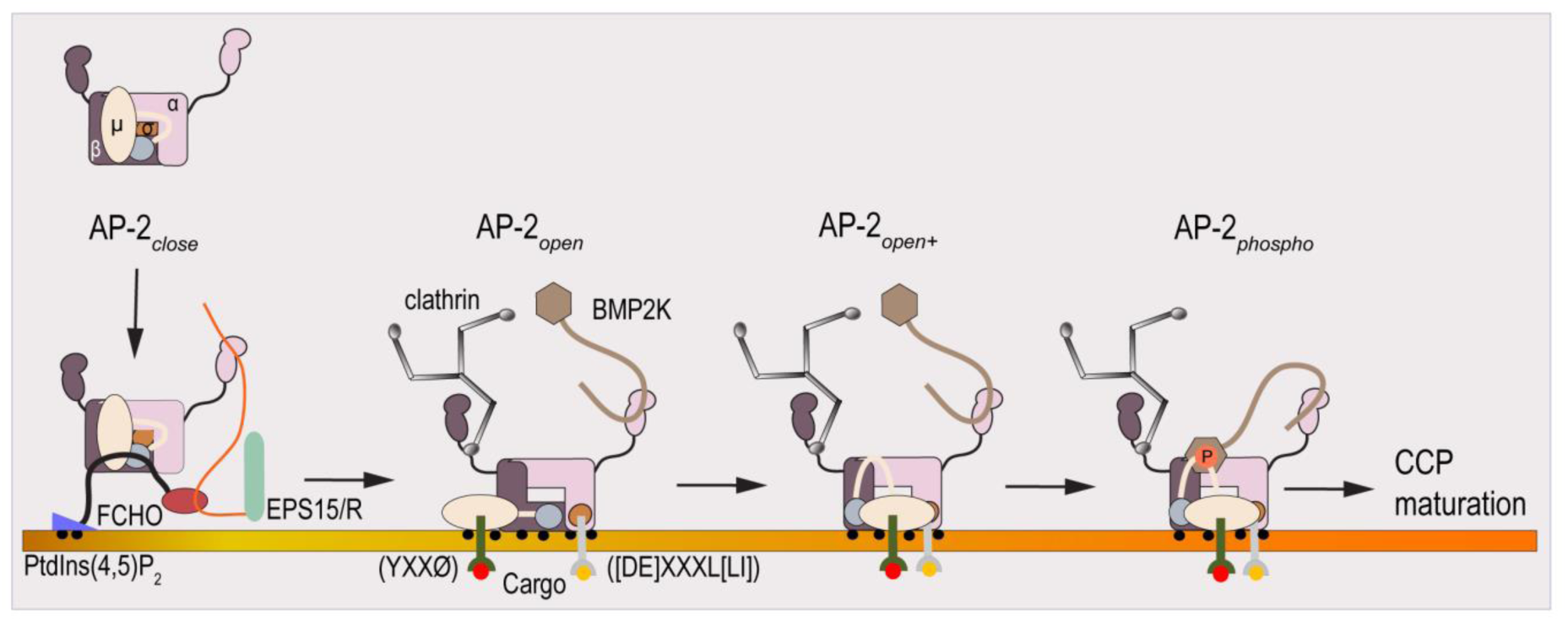
Schematic model depicting role of allosteric FCHO-BMP2K axis in AP-2 activation. FCHO proteins together with EPS15/R allosterically drive the AP-2 equilibrium from cytosolic closed state (AP-2_closed_) to open / open+ conformation (AP-2_open_ / AP-2_open+_) on membrane. AP-2_open_ / AP-2_open+_ recruits and stabilizes BMP2K through appendages without affecting membrane phospholipid [PtdIns(4,5)P2] binding, clathrin binding through β hinge and cargo binding through µ (YXXØ type cargo) and σ ([DE]XXXL[LI] type cargo) subunits as indicated. BMP2K phosphorylates only open+ AP-2 at the exposed T156 residue on µ linker (AP-2_phospho_). AP-2_phospho_ further drives CCP maturation by recruiting late effector proteins.

Degradation of BMP2K in FCHO knockout cells and AP-2 depleted cells reveals unique facets of AP-2 kinases and novel regulatory mechanisms for AP-2 activation. Albeit, it appears that BMP2K can associate with all conformations of AP-2, it is likely prone to ubiquitination and degradation in cytosol and hence has a short half-life in cells (Lu and Hunter, 2009). Extremely low abundance of BMP2K in HeLa cells observed in this study, low copy numbers of kinase assessed by proteomic analyses (Hein et al., 2015; Itzhak et al., 2016) and presence of several possible ubiquitination sites on kinase domain (Akimov et al., 2018) suggest this possibility. Thus, recruitment to CCP could extend the longevity of kinase by protecting it from cytosolic E3 ubiquitin ligases (Piper et al., 2014). CCP recruitment pattern of BMP2K further substantiates this idea. Despite having a long C-terminus (∼ 600 aminoacids) containing an array of SLiMs to co-ordinate multitude of interactions (Sorkin, 2004), BMP2K binds mainly to AP-2 and appears to depend on AP-2 for CCP recruitment and stabilization. Lack of membrane-binding domains (Itoh and De Camilli, 2006; Hurley, 2006) and presence of poly-glutamate (poly-Q) rich region in kinase playing a definitive role in CCP recruitment implies unique trafficking route for BMP2K. Poly-Q stretches are reported in many proteins but to our knowledge BMP2K (and AAK1) is the only endocytic protein identified in this category (Totzeck et al., 2017). Poly-Q tracts are predicted to play role in protein-protein interactions in a context dependent manner (Schaefer et al., 2012) and also in aggregation and accumulation of proteins leading to degenerative diseases (Orr and Zoghbi, 2007). Thus, our results argue a similar function for poly-Q tract in enhancing C-terminus-AP-2 interactions to accumulate and stabilize BMP2K at CCP. Hence, it is logical to conclude that decreasing the amount of membrane active AP-2, either by reducing gross AP-2 levels (AP-2 *RNAi*) or by inhibiting its allosteric activation (FCHO knockout cells), could retain BMP2K in cytosol for degradation. This mode of kinase regulation may have significance in providing spatio-temporal fidelity to AP-2 activation (Ghosh and Kornfeld, 2003a). Thus, together with active cytosolic phosphatases (Ricotta et al., 2008), stringent regulation of kinases could prevent phosphorylation of AP-2 in the cytosol (Wrobel et al., 2019) that may lead to conformational changes unfavourable for subunit assembly (Gulbranson et al., 2019), membrane binding and activation (Ghosh and Kornfeld, 2003b). We assume that most of the BMP2K attributes are redundant to AAK1. It has been reported that clathrin activates AAK1, so could be similar for BMP2K (Jackson et al., 2003). However, C-terminus and SLiMs appear distinct in both kinases; hence recruitment and stabilization mechanisms may vary and are open for future investigations.

Our findings also suggest that steady-state abundance and function of BMP2K is linked to lattice morphology and status of CME in mammalian cells. According to our model, delay in CME in *BMP2K i* cells may account for shift in equilibrium toward AP-2 conformations that can still bind cargo and assemble clathrin but cannot couple downstream effectors of endocytosis *viz*; NECAP (Wrobel et al., 2019; Partlow et al., 2019). This could cause kinetic uncoupling of early and late stages of CCP formation leading to delayed maturation and reduced CME (Nunez et al., 2011; Lampe et al., 2016). Alternatively, accumulation of excess unphosphorylated AP-2 in kinase inactive CCPs could interfere with auxilin/ Hsc70-dependent clathrin exchange during coat assembly leading to aberrant kinetic proofreading controls of endocytosis (Avinoam et al., 2015; Chen et al. 2019). Another possibility is the activation of a compensatory endocytosis mechanism which could bypass sequential AP-2 activation events so as to ensure slow but steady actin-based endocytosis in HeLa cells (Leyton-Puig et al., 2017). Regardless, all these cases would result in deposition of excess AP-2 and immobile clathrin population on membrane giving rise to structures >0.6 µm^2^ as seen in this study as well as in previous studies (Umasankar et al., 2014; Ma et al., 2016; Traub., 2019). Since our current investigation is limited to qualitative analyses of endocytosis phenotypes, these possibilities are speculative and require future investigations to delineate precise mechanisms. However, such effects of kinase and AP-2 phosphorylation are very well consistent with our own results as well as with similar phenotypes observed in *C. elegans* mutants of related NIMA family of kinases (Yochem et al., 2015; Lazetic and Fay, 2017; Lazetic et al., 2018) or in mammalian cells upon depletion of late endocytic effectors (NECAP/ SNX9/ Amphiphysin) (Wrobel et al., 2019). Considering our speculations, BMP2K not correcting lattice morphology defects in FCHO knockout cells would argue AP-2 conformations that are not favourable for phosphorylation in these cells. Alternatively, in FCHO knockout cells, AP-2 may not fully open out and so cannot function regardless of phosphorylation status. Along with supporting these possibilities, reexpression of FCHO correcting defects associated with CCP morphology, AP-2 phosphorylation and function further reveals directionality and sequentiality of allosteric axis.

Though fundamentally similar, modalities of CME are different in invertebrates and vertebrates (McMahon & Boucrot, 2011). 60% of CME proteins have paralogs in vertebrates as compared to 12% in invertebrates (Dergai et al., 2016). Furthermore, AP-2 depletion is viable in *C.elegans* (Gu et al., 2013; Hollopeter et al., 2014), but embryonically lethal in zebrafish (Umasankar et al., 2012 and this study) and mice (Mitsunari et al., 2005). Mechanisms of allosteric regulation of AP-2 and its significance also reflect differently in *C. elegans* and mammals (Hollopeter et al., 2014; Umasankar et al., 2014; Beacham et al., 2018; Partlow et al., 2019; Wrobel et al., 2019). Our *in vivo* studies in zebrafish are consistent with mammalian cell-based results and show FCHO-BMP2K axis co-ordinates AP-2 activation during embryogenesis. Further, human FCHO2 and BMP2K can activate zebrafish AP-2 to drive CME implies allosteric axis is evolutionarily conserved across vertebrates. Perturbing the axis by silencing *fcho2* alone (this study) or together with *fcho1* (Umasankar et al., 2012) seem to affect only a subset of AP-2 functions during embryonic development. This would argue FCHO-BMP2K axis independent *in vivo* activation mechanisms, differential cargo sorting requirements (Kelly et al., 2008) or conformation-independent functions (Gu et al., 2013) for AP-2 during embryogenesis. Thus FCHO and BMP2K appear to have evolved their role in vertebrates as positive modulators of AP-2 function. Hence, we propose that by inducing functionally distinct AP-2 conformations, FCHO-BMP2K axis provides a mechanism to link CCP initiation with CCP maturation to control spatio-temporal mechanodynamics of CME in vertebrates. Since µT156 residue is conserved in other clathrin adaptor protein complexes, it is tempting to speculate similar feed-forward mechanisms for regulation of membrane trafficking events in various cellular compartments.

## Acknowledgements

PKU dedicates this manuscript to Prof. Linton Traub for being a great mentor and for providing endless support and advice. PKU acknowledges Prof. M. Radhakrishna Pillai, Director, Rajiv Gandhi Centre for Biotechnology, for lab facilities and support. PKU is thankful to Prof. Ajay Parida, Director, Institute of Life Sciences for timely help with zebrafish facility. PKU thankfully acknowledges valuable suggestions of Scientific Advisory Council, Rajiv Gandhi Centre for Biotechnology, Prof. Vijayalakshmi Ravindranath and Prof. Balachandran Ravindran in this study. We are extremely grateful to Prof. Linton Traub for graciously providing important reagents for this study. We thank Dr. T. R. Santhoshkumar and Central Imaging Facility, RGCB for help with microscopy. We are grateful to Dr. Sachin Holkar for helping us with quantitation of fluorescence confocal images. We acknowledge technical assistance of Yedukrishnan M., Reshmi P. B. and Merlin Jeejo in the early stages of this work and also thank Anup Biswal for maintaining zebrafish facility.

## Materials and Methods

### Molecular cloning and constructs

Sequences encoding human BMP2K (1-1161), BMP2K (1-560) and BMP2K (561-1161) were PCR amplified from the template ORF clone # RC215795 (Origene) and inserted into *HindIII* restriction site of pEGFP-N1 vector using In-Fusion cloning technology (Clontech) to generate BMP2K-GFP, BMP2KΔCT-GFP and BMP2KCT-GFP respectively. Cloning of PCR amplified BMP2K (1-345) or BMP2K (346-560) into an upstream in-frame *NheI* site in the BMP2KCT-GFP construct yielded BMP2KΔQ/H-GFP or BMP2KΔKD-GFP respectively. GFP-FCHO1 and GFP-FCHO1 (1-609) were designed as described previously (Umasankar *et al*., 2014). Constructs encoding BMP2K regions fused to N-terminal GST *viz*; GST-BMP2K-KD (38-345), GST-BMP2K-Q/H (346-560), GST-BMP2K-CT#1 (561-850) and GST-BMP2K CT#2 (851-1161) were generated by the insertion of corresponding PCR amplicons between *EcoRI* and *NotI* restriction sites in pGEX-4T-1 vector through directional cloning. GST-rat β2 appendage (701–937), GST-mouse α_C_ appendage (701–938) and GST-human EPS15 (595–896) have been described previously (Umasankar *et al*., 2012). pCS2-*Hs* BMP2K and pCS2-*Hs* FCHO2 were generated by cloning human BMP2K (1-1161) and human FCHO2 (1-810) (Dharmacon) coding sequences respectively into the *EcoRI* site of pCS2^+^ vector using In-Fusion cloning. Introducing stop codon at residue 561 in pCS2-*Hs* BMP2K plasmid using QuikChange site-directed mutagenesis (Agilent Technologies) yielded pCS2-*Hs* BMP2KΔCT. All constructs were verified by automated dideoxynucleotide sequencing (Eurofins). Complete details of sequences and plasmid maps are available upon request.

### Cell culture and transfections

The authenticated and validated parental HeLa SS6 cell line (Umasankar *et al*., 2014, Traub 2019) was a kind gift from Prof. Linton Traub. HeLa clone #64/1.E (1.E cells) was generated previously by bi-allelic disruption of the FCHO2 and FCHO1 gene loci using TALEN gene-editing technology (Umasankar *et al*., 2014). HeLa SS6 and 1.E cells were cultured in complete medium-Dulbecco’s modified Eagle’s medium (DMEM, Hyclone) supplemented with 10% fetal bovine serum (FBS, Gibco) and 2 mM L-glutamine (Gibco) and were maintained at 37°C in a CO_2_ incubator (Thermo scientific) under 5% CO_2_. Cells were passaged at 90%-100% confluence and were periodically checked for mycoplasma contamination using commercially available kits.

For all transfections, HeLa SS6 or 1.E cells were seeded into 35 mm tissue culture dishes or 6 well tissue culture plates (Eppendorf) with or without pre-cleaned, 12 mm round sterile glass coverslips, one day before transfection and were grown in complete medium at 37^0^C under 5% CO_2_. All transfections were done when cells reached 50%-60% confluence. Transfected cells grown on coverslips were used for endocytosis assays and immunofluorescence, whereas those in dishes without coverslips were lysed for immunoblotting. Endotoxin-free miniprep or midiprep plasmid DNA transfections were performed using Lipofectamine 2000 (Invitrogen). On the day of transfection, 6 µl Lipofectamine 2000 was added to 0.5-1.0 µg plasmid DNA diluted in 375 µl Opti-MEM I Reduced Serum medium (Gibco) to make a transfection mixture and incubated for 40 minutes at room temperature (RT). This transfection mixture was added drop wise to cells replenished with 1.5 ml fresh complete medium. Cells were further grown at 37^0^C, 5% CO_2_ for 18-24 hours post-transfection, and were used for various experiments.

### RNAi and inhibitor treatments

RNAi experiments were performed by knocking down the gene targets in HeLa SS6 cells growing in 35 mm cell culture plates with two rounds of siRNA transfections using Oligofectamine (Invitrogen) as described (Motley *et al*., 2003). Custom synthesized siRNA (10 nmol) and siGENOME SMARTPOOL siRNA (5 nmol) oligonucleotides (Key Resources Table) were obtained from Dharmacon and resuspended in 1X siRNA buffer (Dharmacon) to make 20 µM stock. On Day-1, 10 µl siRNA stock solution was diluted in 175 µl Opti-MEM I in a sterile microfuge tube. 5 µl Oligofectamine was diluted in 10 µl Opti-MEM I in a separate tube. After incubating for 10 minutes at RT, solutions in two tubes were mixed and incubated for another 20 minutes at RT for transfection-complex formation. Meanwhile, 800 µl Opti-MEM I was added into each dish after removing complete medium. 200 µl transfection mix was added drop wise into each dish and cells were incubated for 4-6 hours post-transfection in serum free-medium. Then, 500 µl opti-MEM I containing 30% FBS was added into cells and incubated for another 18-24 hours at 37^0^C in 5% CO2 atmosphere. On Day-2, transfected cells were trypsinized and seeded into new dishes in duplicates or triplicates for second round of knockdown. On Day-3, second round of siRNA transfections were performed in the same fashion. On Day-4, cells were ready for various experiments. Co-transfection of plasmid DNA was performed along with second round of siRNA transfections using Lipofectamine 2000 instead of Oligofectamine. All RNAi experiments were repeated at least three times and each experiment included control dishes with mock transfected cells that received all reagents except siRNA. Each set of experiments were analyzed for knockdown efficiency by comparing the levels of target gene expression in mock and RNAi samples through immunoblotting.

Calyculin (Sigma) and SGC-AAK1-1 (Tocris) inhibitors dissolved in DMSO were used for various experiments as mentioned in the text. HeLa SS6 cells in culture dishes received a final concentration of 50 nM calyculin for 1 hour or 10 µM SGC-AAK1-1- for 2 hours in complete medium at 37^0^C, 5% CO_2_. Control dishes with mock treated cells were included in all experiments that received same concentration of DMSO for the same time period as cells in experimental dishes. Calyculin treatment caused cells to round up and detach from the substratum. After treatment, cells were stripped from dishes using Non-enzymatic Cell Dissociation Solution (Sigma) at RT and collected into microfuge tubes on ice. PBS containing calyculin or SGC-AAK1-1 or DMSO was used to rinse corresponding dishes and pooled with Cell Dissociation solution in respective tubes. Cells were pelleted by centrifuging at 500 × *g_max_* for 5 minutes at 4^0^C. After discarding the supernatant, cell pellets were resuspended in pre-heated SDS-Laemmli buffer. Alternatively, cells grown on coverslips, incubated for one hour with complete medium followed by another hour with starvation medium, both containing appropriate concentrations of SGC-AAK1-1 or DMSO were used directly for endocytosis assays.

### Endocytosis assays

Endocytosis assays were performed using three different protocols. In all protocols, cells growing on coverslips in complete medium (DMEM, 10% FBS, 2mM L-glutamine) at 37^0^C in 5% CO2 atmosphere were first starved by incubating in starvation medium (DMEM, 25mM HEPES, pH-7.2, 0.5% BSA, 2mM L-glutamine) for 1 hour. This will undock bound apo-transferrin from cell surface transferrin receptors thereby making the receptors available for fluorescent transferrin binding. After removing the starvation medium, cells were subsequently pulsed at 37^0^C for different time points with 25µg/ml Alexa flour-488 or- 568-conjugated human transferrin (Invitrogen, ThermoFisher Scientific) reconstituted in starvation medium and clarified by centrifugation at 15,000 × g_max_ at 4^0^C for 10 minutes. In protocol #1, after a 2 minute or 10 minute pulse at 37^0^C, cells were immediately transferred on ice, washed three times with ice-cold PBS and fixed with 2% paraformaldehyde (Electron Microscopy Sciences) for 20 minutes at RT. In protocol #2, after 5 minute pulse, cells were washed three times with stripping buffer (0.2 M acetic acid, 0.2 M sodium chloride, pH 2.5) on ice to remove surface-bound pool of transferrin and fixed with paraformaldehyde. In protocol #3, a modified endocytosis assay was used. In this assay, after 2 minute pulse of fluorescent transferrin at 37^0^C, cells were surface stripped with stripping buffer on ice as in protocol #2. Then, cells were rewarmed to 37^0^C for another 2 minutes to continue endocytic transport before paraformaldehyde fixation (Reis et al., 2015; Ma et al., 2016).

### Immunofluorescence analysis

Immunofluorescence analysis was performed as described (Umasankar et al., 2014; Ma et al., 2016). Paraformaldehyde fixed cells on 12 mm round glass cover slips were washed three times in ice-cold PBS. Then, cells were blocked and permeabilized by incubating in 10% normal goat serum (Himedia), 0.2% saponin (Sigma) in PBS at RT for 30 minutes. Primary antibodies at appropriate concentrations were diluted in antibody dilution buffer (10% normal goat serum, 0.05% saponin in PBS) and centrifuged at15,000 × *g_max_* for 10 minutes before adding into permeabilized cells. Primary antibody incubation was for 1 hour at RT followed by three washes with PBS. Anti-mouse/ rabbit secondary antibodies conjugated with the fluorophores-Alexafluor 488/568/647 (Invitrogen, ThermoFisher Scientific and Cell Signaling Technologies) were diluted 1:500 in antibody dilution buffer and were clarified by centrifugation. After secondary antibody incubation for one hour at RT in the dark, cells were washed three times with ice-cold PBS. Cells were further incubated with 1 µg/ml Hoechst 33342 DNA dye (Invitrogen, ThermoFisher Scientific) for 10 minutes at RT and washed twice with ice-cold PBS before mounting onto glass slides with 5 µl Fluoromount-G mounting medium (Electron Microscopy Sciences).

### Confocal Microscopy and Image Processing

Images were acquired using Nikon A1R Eclipse Ti Confocal Laser Scanning Microscope equipped with Plan Apo VC 60x oil objective, NA-1.4. Different lasers were used to excite fluorophores at various wavelengths – 405 nm laser for Hoechst, 488 nm laser for Alexa Fluor 488, the 561 nm laser for Alexa Fluor 568 and 638 nm laser for Alexa Fluor 647 as programmed in the Nikon NIS Elements software. Emission signals from different channels were collected as 1024 × 1024 pixel images in sequential scan mode with line averaging function. Z-stacks were obtained by sequential scans of 0.3 µm step size between each optical section and were merged to obtain max-Z projection. All images were saved in nd2 format and later converted to TIFF files using NIS Elements software. All images were equally cropped, adjusted and processed in Adobe Photoshop CS3 for better visualization, arranged and labelled in Adobe Illustrator CS6.

### Quantitation of confocal micrographs

Quantitation of fluorescence confocal images was done as described previously (Ma et al., 2016) using Nikon NIS Elements AR 3.2 analysis package and Fiji (ImageJ). Before analysis, confocal optical sections of single plane or Z-stacks for AP-2, pT156µ or transferrin channels were deconvoluted using AutoQuant X3-2D/ 3D deconvolution software and merged into single maximum projection images. Images of individual cells to be analyzed were dissected out from the whole field by creating region-of-interests (ROI), converted to binary images by thresholding and background subtraction. Single cell masks were generated from binay images and were used to analyze size (area) or fluorescent intensity of objects in the background corrected raw images for corresponding original channels. For CCP morphology analysis, objects were binned into area-based categories as described previously (Ma et al., 2016) and as explained in the figure legends. Area in pixels was converted to µm^2^ (1 pixel= 0.1 µm) and used in all calculations. In all analysis, total cellular areas were measured from ROIs to determine density and mean intensity per µm^2^. Mostly, images from 3 independent experiments were separately analysed before compiling the data. Data was compiled and analyzed in Microsoft Excel and graphs were plotted in GraphPad Prism.

### Recombinant protein expression and purification

Control GST and GST-fusion proteins were expressed through pGEX-plasmids in *E.coli* BL21 codon plus RIL strains and purified using standard procedure (Umasankar et al., 2012; Umasankar et al., 2014; Ma et al., 2016). Large volumes of 2x YT media (Sigma) inoculated with overnight grown bacterial culture was grown at 37^0^C till absorbance A_600_ reaches ∼0.6 and induced with 100 µM IPTG for 4 hours at 22^0^C. Bacterial pellets were collected by centrifugation at 15,000 × g_max_ at 4^0^C for 10 minutes and frozen at −80^0^C. Frozen pellets were resuspended in ice-cold bacterial lysis buffer (50 mM Tris-HCl, pH-8.0, 300 mM sodium chloride, 10 mM β-mercaptoethanol and 0.2% (v/v) Triton X-100) containing 1 mM PMSF (Sigma) on ice. After disrupting cells with five pulses of sonication using a Vibracell microprobe sonicator (Sonics) at an interval of 1 minute between each pulse, the lysate was centrifuged at 15,600 × g_max_ at 4^0^C for 30 minutes. The clarified supernatant was mixed with Glutathione Sepharose (GE Life Sciences) beads and incubated with continuous mixing for 2 hours at 4^0^C on a tube nutator. Beads were collected at 500 × g_max_ at 4^0^C and washed four times with ice-cold PBS. Bound protein was eluted three times in Elution Buffer-25 mM Tris-HCl, 250 mM sodium chloride, 10 mM reduced glutathione (Sigma) and 2 mM DTT on ice. After assessing purity and yield of eluted proteins on SDS-PAGE, the elutions were pooled and dialysed overnight in PBS. Final protein concentrations were determined using Bradford method with BSA as standard. All purified proteins were aliquoted and stored at −80^0^C until further use.

### GST pull-down assays and Immunoblotting

HeLa SS6 cell lysate (prey) for GST-pull down assays were made as described (Umasankar et al., 2012; Umasankar et al., 2014; Ma et al., 2016). Briefly, HeLa SS6 or 1.E cells grown in several 10 cm tissue culture plates were trypsinized at confluence and pooled into 50 ml polypropylene tubes. Cell pellets were recovered by centrifugation at 1000 × g_max_ at 4^0^C for 10 minutes, washed once with PBS and directly resuspended in ice-cold Cell Lysis Buffer-25 mM HEPES-KOH, pH 7.2, 125 mM potassium acetate, 5 mM magnesium acetate, 2 mM EDTA, 2 mM EGTA, 2 mM DTT and 1% Triton X-100 containing cOmplete EDTA-free Protease Inhibitor Cocktail on ice. The suspension was incubated on ice for 30-45 minutes with occasional mixing and sonicated for 10 seconds. After centrifugation at 20,000 × g_max_ for 30 minutes to remove insoluble material, the clarified cell lysate was stored in frozen aliquots at −80^0^C.

In pull-down assays, appropriate concentrations of GST or GST-fusion protein was immobilized on 30 µl Glutathione Sepharose beads (bait) in microfuge tubes containing 750 µl PBS by continuous up-down mixing at 4^0^C. After 2 hours mixing, the beads were recovered at 10,000 × g_max_ for 1 minute and were washed twice with ice-cold assay buffer (25 mM HEPES-KOH, pH 7.2, 125 mM potassium acetate, 5 mM magnesium acetate, 2 mM EDTA, 2 mM EGTA) containing 2 mM DTT. After second wash, supernatant was completely removed and beads were resuspended in 250 µl clarified cell lysate (prey) diluted in assay buffer. Beads were further incubated for 1 hour at 4^0^C with up-down mixing and recovered by centrifugation at 10,000 × g_max_ for 1 minute at 4^0^C. After storing 60 µl (supernatant fraction), rest of the supernatant was carefully discarded. The assay was terminated by washing the beads (pellet fraction) twice with ice-cold PBS containing 0.2 % Triton-X-100, followed by two washes with only PBS. For pull down assays with AP-2 core, a modified assay buffer (assay buffer with 25 mM Tris-HCl, pH-7.2, 0.2% Igepal CA-630 and 10 mM DTT) was used. 250 µl clarified prey containing 25 µg/ ml purified AP-2 core and 0.1 mg/ml BSA in modified assay buffer was added to the immobilized bait. Supernatant and pellet fractions were recovered in a similar fashion. After final wash with PBS, SDS-Laemmli buffer was added to both pellet and supernatant fractions to make a final volume of 80 µl. Supernatant and pellet fractions were resolved on SDS-PAGE and gels were either stained with Coomassie blue or transferred to nitrocellulose membrane for immunoblotting. Blots were blocked in 5 % non-fat dried milk powder in TBST at RT, and then incubated with primary antibodies diluted in 1% milk-TBST for 1-2 hours at RT. After washing with TBST, blots were incubated with 1: 5000 dilution of HRP-conjugated anti-mouse/ rabbit IgG secondary antibodies (Bio-Rad) in 1% milk-TBST for another hour. Blots were then washed in TBST and the bands were developed with Amersham ECL (GE Life Sciences) or Clarity ECL (Bio-Rad) western blotting substrates using autoradiography.

### Zebrafish maintenance and husbandry

*Danio rerio* AB strains were maintained under standard conditions in accordance with guidelines for use, care and maintenance of experimental animal models and with Institutional Animal Ethical Committee (IAEC) approval. Typically, male and female fishes were grown in multilinked aquatic units (Techniplast) equipped with circulating water at 28.5^0^C and at pH 7.5. Photoperiodicity was set with 14 hours light, 10 hours dark cycle; fishes were provided standard fish food. Male and female fishes from the same batch were separated overnight in breeding tanks and were mixed together to mate and spawn on the day of experiment. Fertilized embryos from natural matings were obtained in batches and were developmentally staged for various experiments. Atleast two weeks of interval was given in between each breeding cycle to avoid stress to same batch of male and female fishes.

### *In vitro* transcription, morpholinos and microinjections

pCS2+ vertebrate expression vectors containing human-BMP2K, BMP2K ΔCT or FCHO2 coding sequences were used as templates to synthesize capped mRNAs for ectopic expression and rescue experiments. Capped mRNAs were synthesized by in vitro transcription reaction from linearized pCS2+ constructs using SP6 mMessage mMachine kit (Ambion) by following vendor’s instructions. Each in vitro transcribed mRNA was extracted with equal volume of phenol/ chloroform and precipitated with chilled isopropanol. Precipitated RNA pellet was dissolved in 10-20 µl nuclease free water (Ambion). After spectrophotometric estimation of concentration, each capped mRNA was stored as aliquots at −800C until used for microinjections.

Translation blocking ATG specific morpholinos were custom synthesized by Gene tools to silence *D. rerio fcho2* and *ap-2a1* genes (Key Resource Table) through anti-sense technology as described (Umasankar et al., 2012). Standard control morpholino oligonucleotide was also obtained from Genetools and was used as injection control in various experiments. Lyophilized morpholinos obtained from Genetools were reconstituted in 300 µl nuclease free water to obtain 1 mM final stock solutions and were stored as frozen aliquots.

*D. rerio* embryos at one-cell stage were used for all microinjections. Typically, 1 nl of morpholino or mRNA diluted in nuclease-free water at desired concentrations were microinjected into the yolk along with 1% phenol red dye using FemtoJet microinjector (Eppendorf). Microinjections for rescue experiments were performed as two separate sets with initial mRNA injections followed by morpholino injections in the same embryo. Each experiment was repeated in three separate biological replicates to get statistically significant numbers to validate our observations. Each time, embryos from the same clutch were used as controls for comparing morphological and molecular parameters. After injections, the embryos were incubated in E3 embryo medium (5mM NaCl, 0.17mM KCl, 0.33mM CaCl2, 0.33mM MgSO4, 0.01% methylene blue) at 28.5^0^C until the desired stage. Unfertilized embryos from both control and experimental samples were identified by microscopic analysis and were removed in all experiments.

### *In vivo* pharmacological experiments

Since pharmacological manipulations in the maternal stages are lethal to developing embryos, we used embryos after ∼ 6 hpf (shield stage) for inhibitor treatments. Embryos from the same clutch were collected and distributed into 24 well plates ensuring that each well receives equal number (ten embryos per well) and that biological samples in triplicates are available for each concentration of inhibitor tested. Triplicates of control samples treated with DMSO were included for each concentration of inhibitor in all experiments. 50 mM stock of SGC-AAK1-1 (AAK1/ BMP2K inhibitor, Tocris) prepared in DMSO was diluted in E3 medium and was used for all experiments. Initially, a pilot assay was performed to normalize the exact concentration of SGC-AAK1-1 to be used for all the experiments. Briefly, embryos were treated with 1, 2, 5, 10, 25 and 50 µM concentrations of SGC-AAK1-1 from 6 hpf to 16 hpf and were screened for convergent extension (CE) defects by morphological analysis. 10 µM concentration of the drug showed specific CE phenotype in all embryos tested and concentrations above showed high precipitation of SGC-AAK1-1 in E3 medium. Hence, a concentration of 10 µM was selected for all experiments. Control embryos and embryos treated with SGC-AAK1-1 were subjected to whole-mount *in situ* hybridization with molecular markers to confirm CE defects.

### Riboprobe synthesis and whole mount *in situ* hybridization (WISH)

Total RNA was isolated from maternal and zygotic stages of *D. rerio* embryos using TRIzol reagent (ThermoFisher Scientific). Single strand cDNA was synthesized from the mRNA fraction of the total RNA by using Oligo dT primer and Superscript III First-Strand Synthesis System for RT-PCR (Invitrogen, ThermoFisher Scientific) according to manufacturer’s instructions. This cDNA was used as templates to PCR amplify various genes expressed in *D. rerio* embryos. PCR amplification using gene-specific primers for *D*. *rerio bmp2k*, *ap2a1* and *dusp6* resulted in amplicons of ∼1 kb size. These amplicons were cloned into pCR Blunt II-TOPO vector. After sequence verification of the clones, plasmids were linearized using compatible restriction enzymes and directly used for riboprobe synthesis. Plasmids containing genes for *no tail* and *myo D* were generously provided by Michael Tsang. Riboprobe synthesis for WISH was carried out with the linearized plasmid using digoxigenin (DIG) RNA labelling mix and T7 RNA polymerase (Roche) according to manufacturer’s recommendations. WISH was performed by standard procedures as described (Umasankar et al., 2012). Briefly, embryos at appropriate developmental stages, with or without chorion, were fixed in 4 % paraformaldehyde for 30 minutes at RT and then at 4^0^C overnight, washed with PBS and stored in methanol at −20^0^C. The embryos were brought back to PBS-Tween 20, treated with Proteinase-K and incubated in pre-hybridization buffer for 2-4 hrs. The embryos were hybridized with respective DIG-labelled antisense probes overnight at 65^0^C. The embryos were washed extensively, blocked with blocking reagent (Roche/Merk) and incubated with anti-DIG antibody conjugated with alkaline phosphatase (AP). BM purple (Roche/Merck) was used as a chromogenic substrate of AP for detecting the gene expression. The reaction was stopped by washing the embryos in PBS several times. Finally, the embryos were mounted in glycerol for imaging.

### Embryonic cell culture

*D. rerio* embryonic cell culture was carried out as described (Westerfield M, 2000). Embryos at ∼ 5 hpf were collected and decontaminated by treating with 0.05% bleach in E3 medium. After de-chorionation with 2 mg/ml Pronase (Sigma) solution in E3, embryos were rinsed in sterile calcium-free Ringer’s solution (116 mM NaCl, 2.9 mM KCl, 5 mM HEPES, pH −7.2). 50 embryos were pooled in a tissue culture plate and were smashed to dissociate embryonic cells by dropping a sterile coverslip on them. The dissociated embryonic cells were pooled into a microfuge tube in sterile calcium-free Ringer’s solution and the suspension was centrifuged at 300 × g_max_ for 7 minutes. After removing supernatant, the pellet was further triturated through a sterile narrow-bore Pasteur pipette to resuspend the cells. Embryonic cells were washed twice with embryonic cell growth medium-L-15 (Sigma), 0.3 mg/ml L-glutamine, 50U/ml penicillin, 0.05 mg/ml streptomycin and 0.8 mM CaCl2 and resuspended in the same medium containing 10% embryo extract and 3% fetal bovine serum. Embryonic cells were plated on sterile glass coverslips coated with 10 µg/ ml laminin (Corning) and grown in six well plates at 28^0^C, without CO_2_ atmosphere.

### *In vivo* imaging of embryos

For live *in vivo* imaging, *D. rerio* embryos with or without chorion were mounted in 3 % methylcellulose (Sigma) in a small petri dish. Embryos at 30 hpf were temporarily anaesthetized by overlaying with E3 containing 0.016 % tricaine to prevent movement during imaging. After orienting laterally, embryos were imaged on Leica MZ16 stereo microscope using Leica LAS X software. For WISH imaging, fixed embryos stained with chromogenic AP substrates were mounted on glycerol in cavity slides and oriented to obtain lateral view. The images were captured with Leica MZ16 stereo microscope. Embryonic cells growing on laminin-coated coverslips were imaged live on a Leica fluorescent microscope using 10x and 20x objectives in phase contrast mode.

### Key Resources table

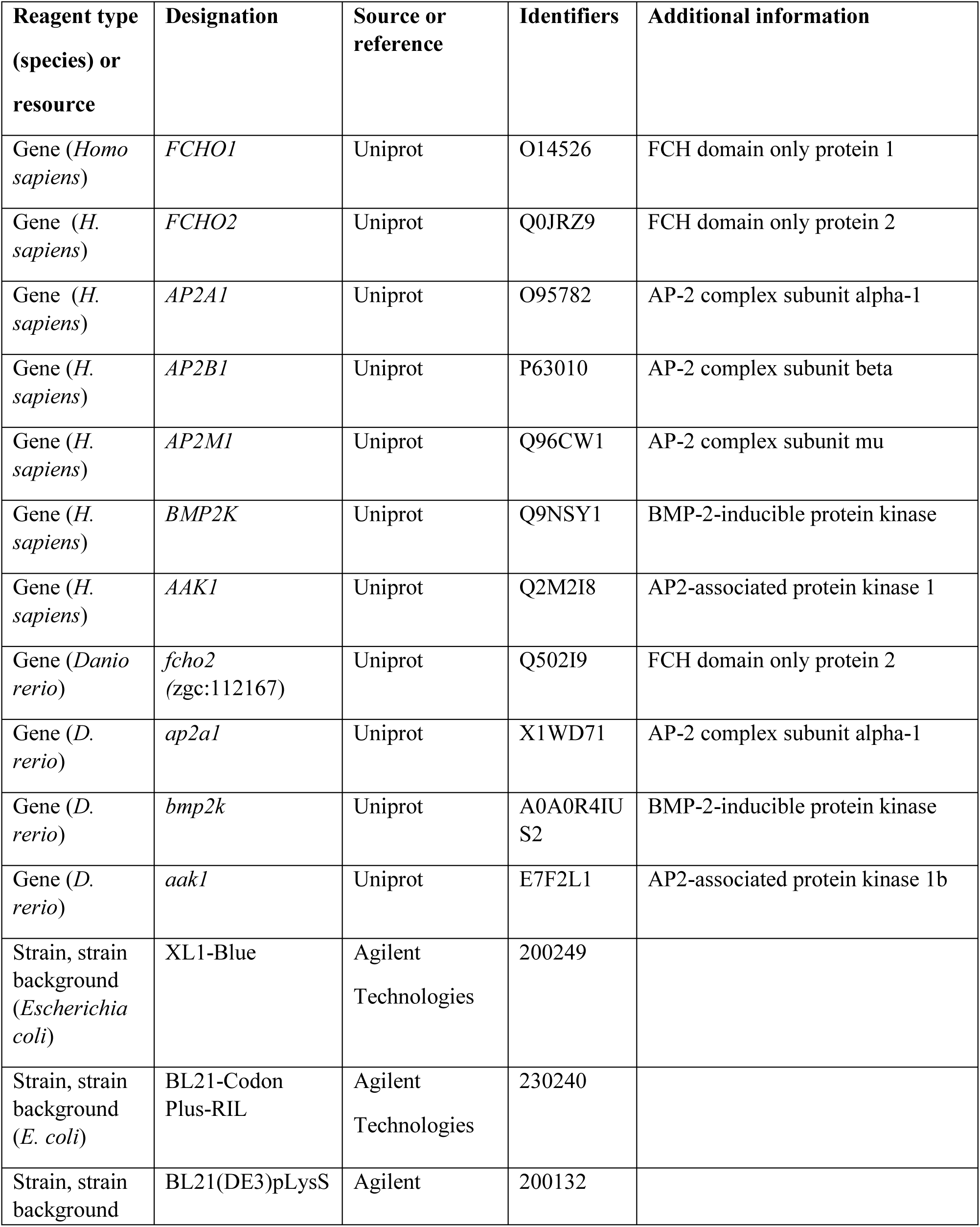
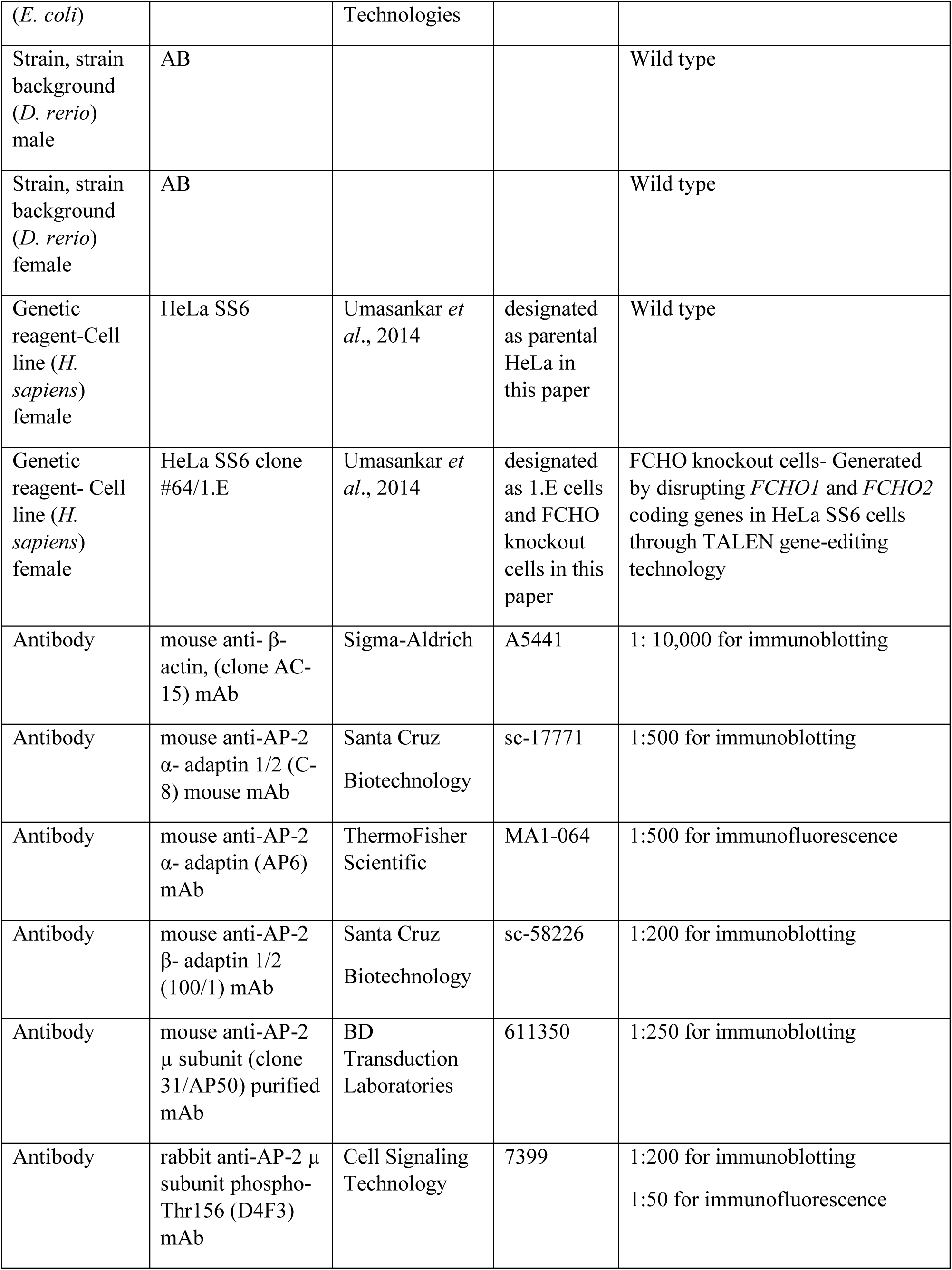
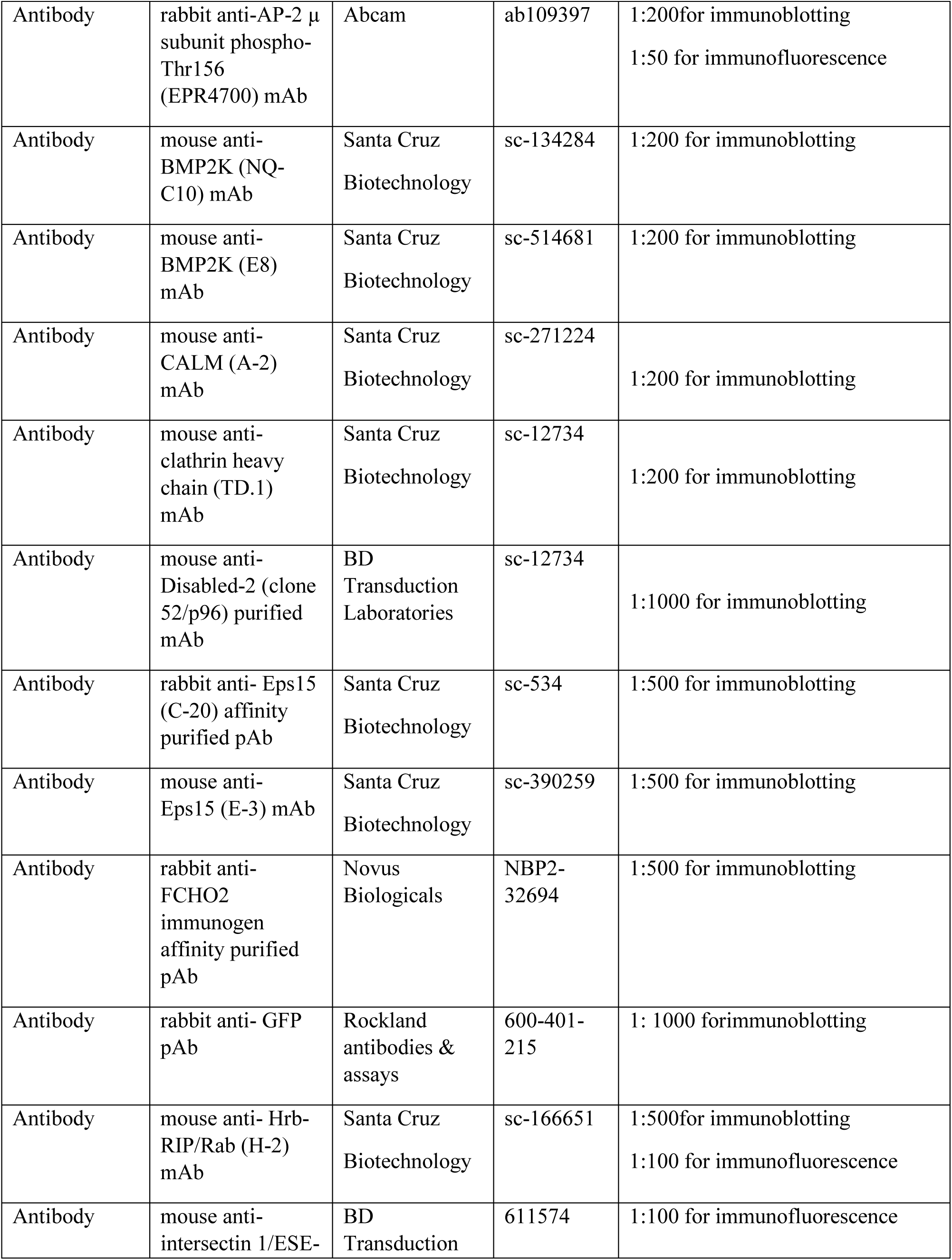
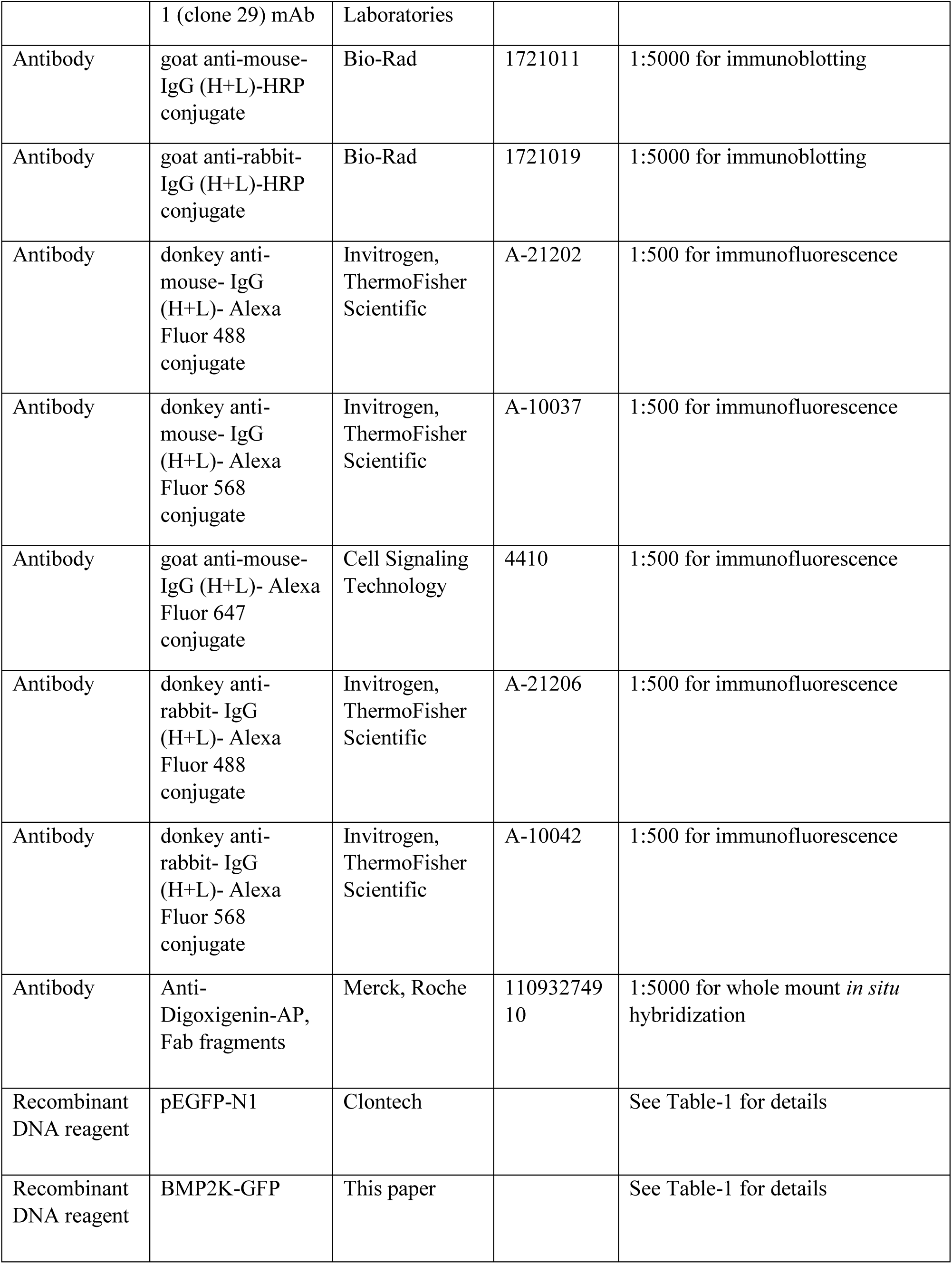
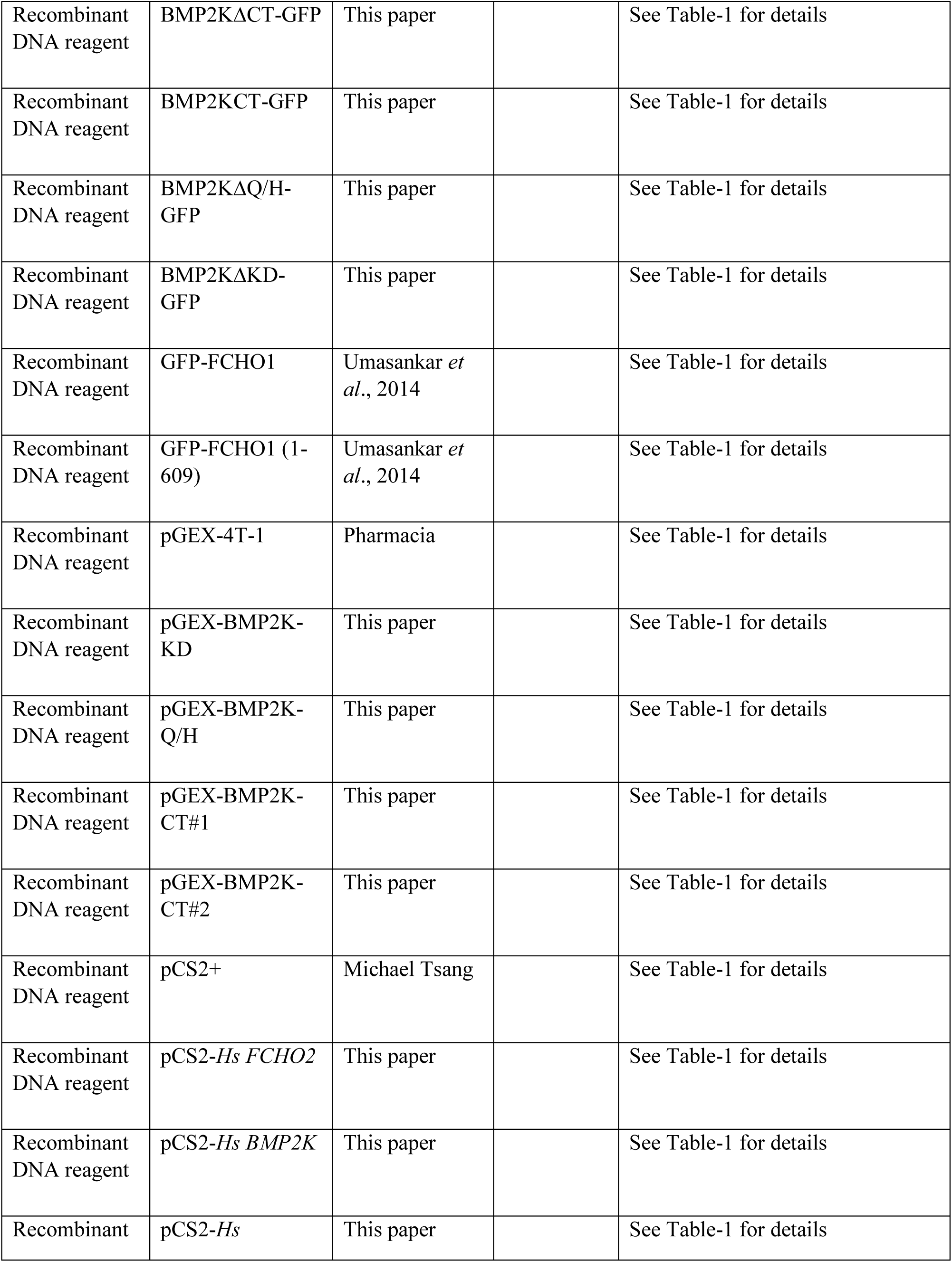
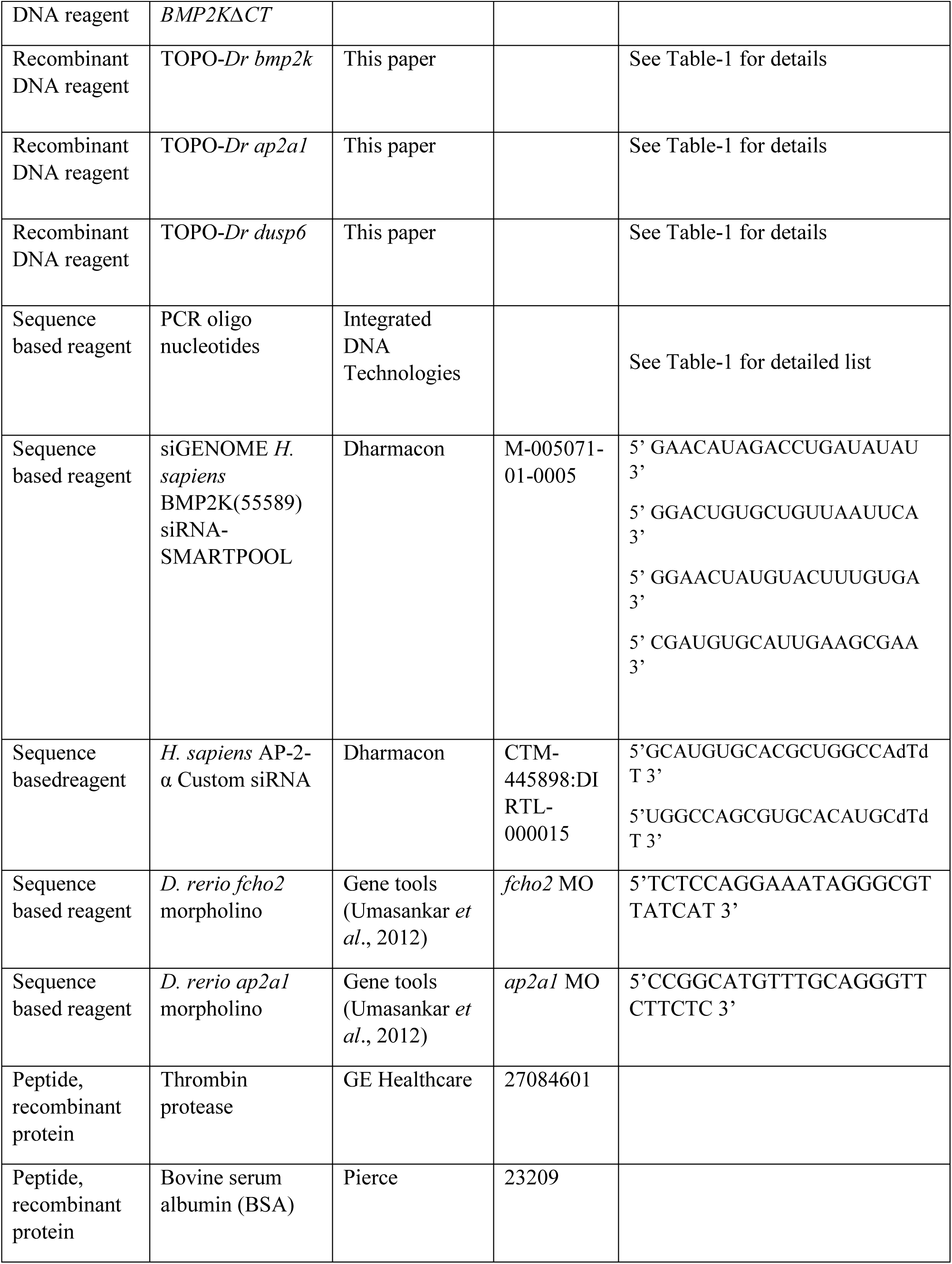
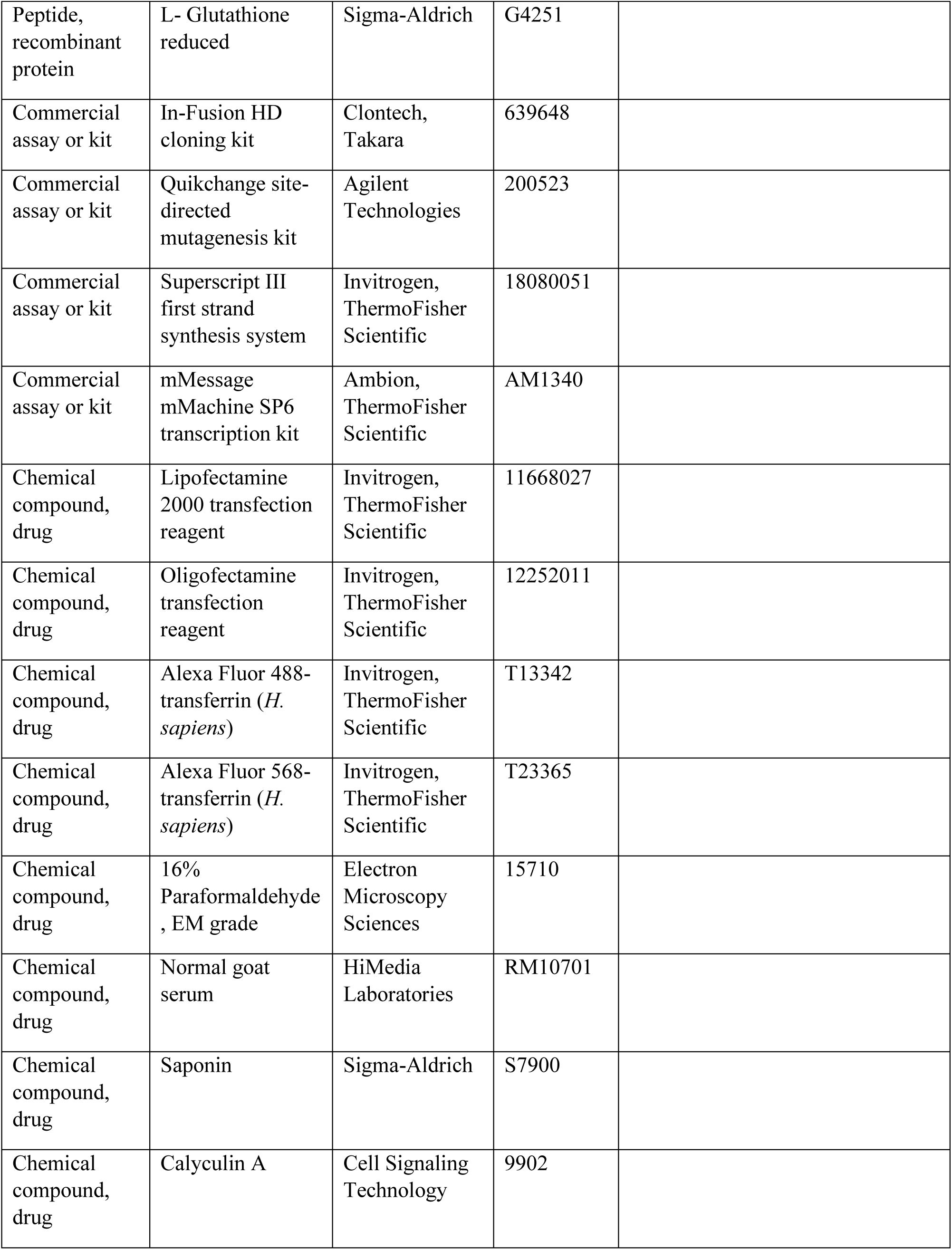
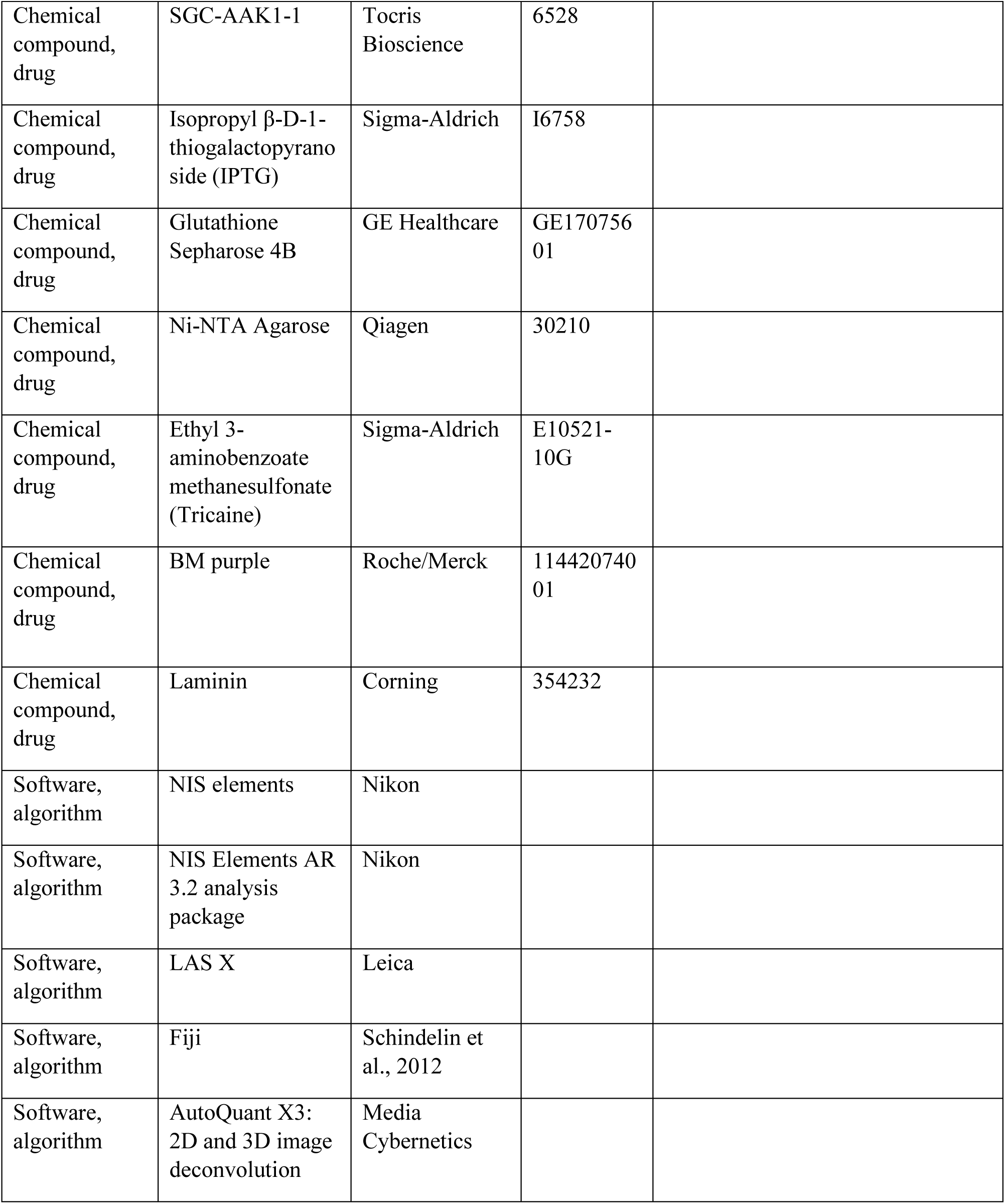

**Table-1:**
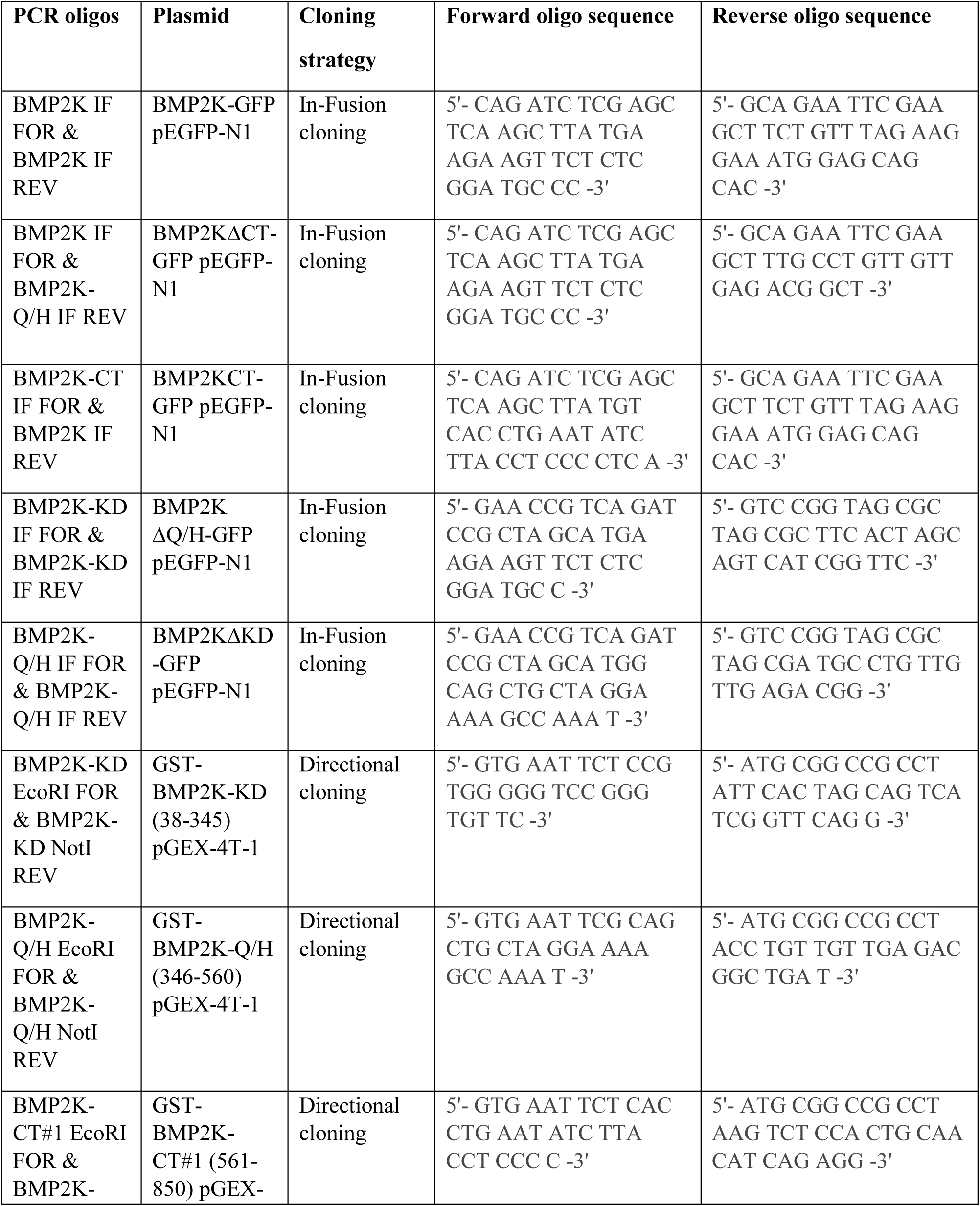
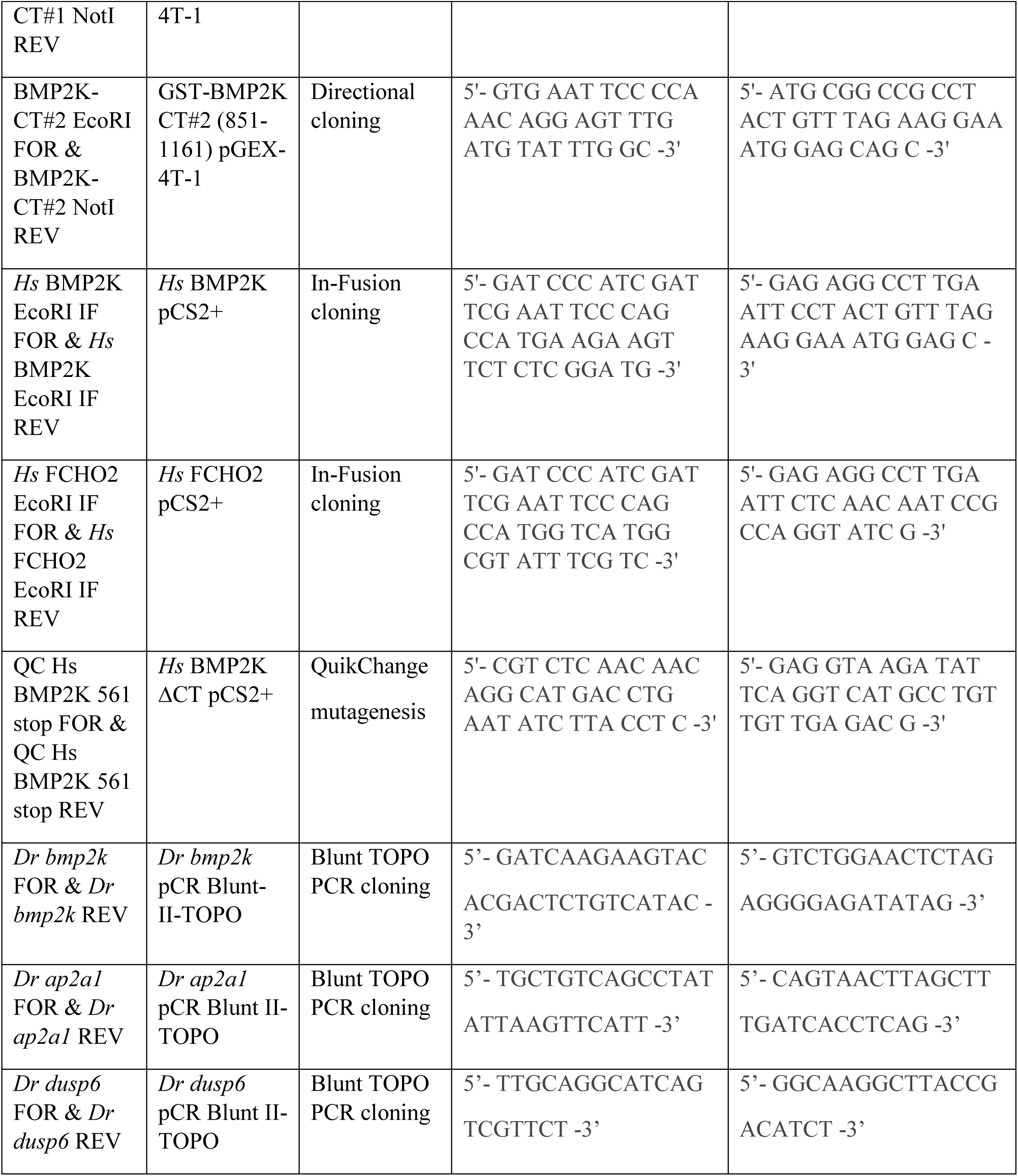
PCR oligonucleotides

---

**Figure 2- figure supplement 1:**
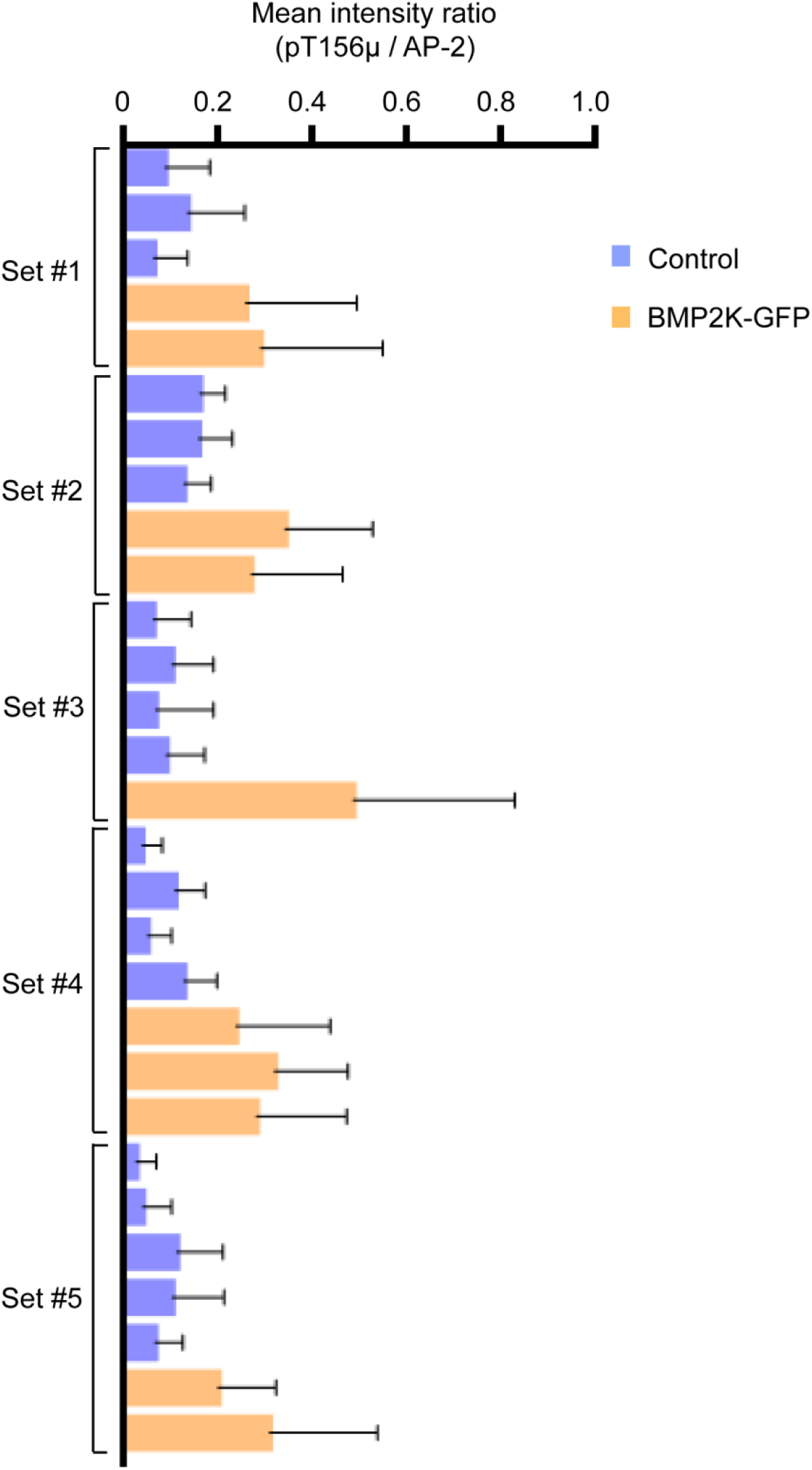
Phosphorylation of AP-2 increases in BMP2K overexpressed cells. Quantification of ratios of mean fluorescence intensities of phosphorylated AP-2 (pT156µ) to total AP-2 in CCPs. Graphs show mean SD of individual BMP2K-GFP transfected (BMP2K-GFP) or untransfected (Control) HeLa SS6 cells randomly picked from 5 confocal data sets.

**Figure 3- figure supplement 1:**
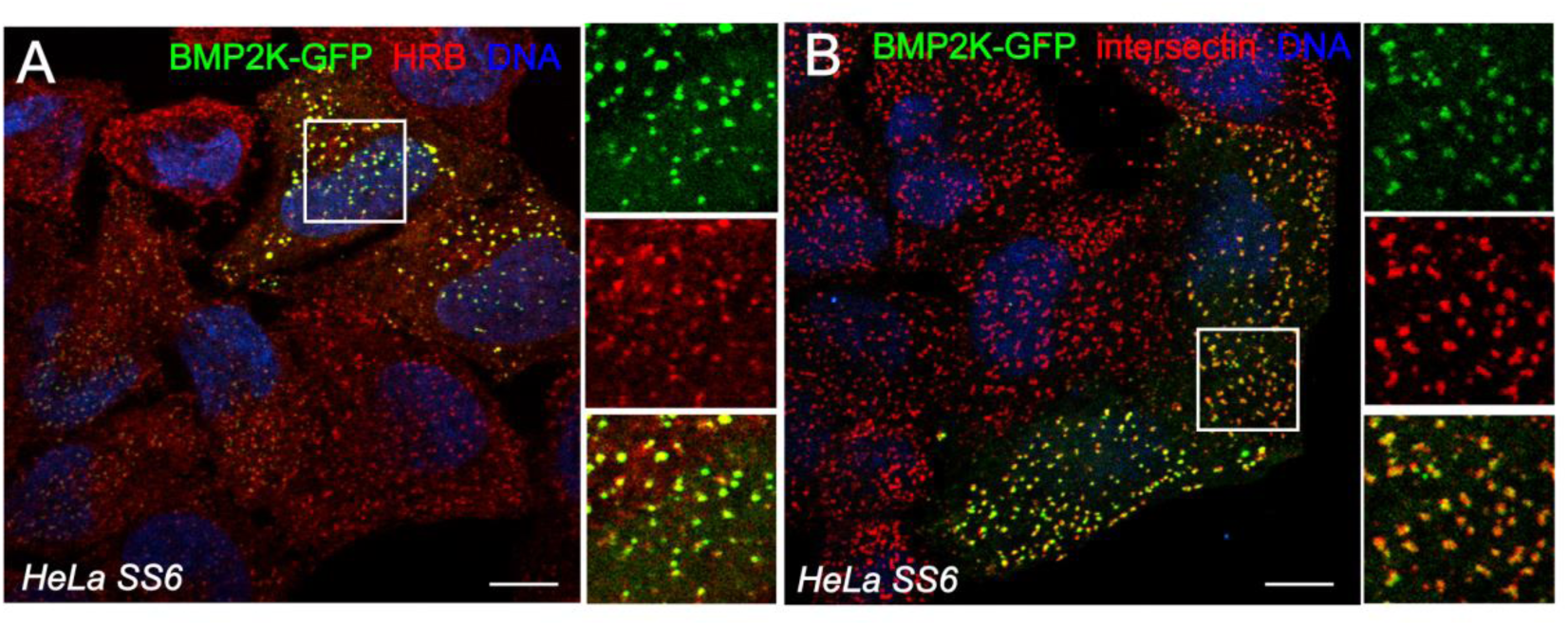
BMP2K co-localizes with CME proteins. (**A and B**) Representative confocal optical sections of the basal plane of HeLa SS6 cells transfected with BMP2K-GFP (green), fixed and immunolabeled for the endocytic proteins, HRB (red) (**A**) or intersectin (**B**). Colour separated channels of the boxed regions are shown on the right. Scale bar-10 µm.

**Figure 6- figure supplement 1:**
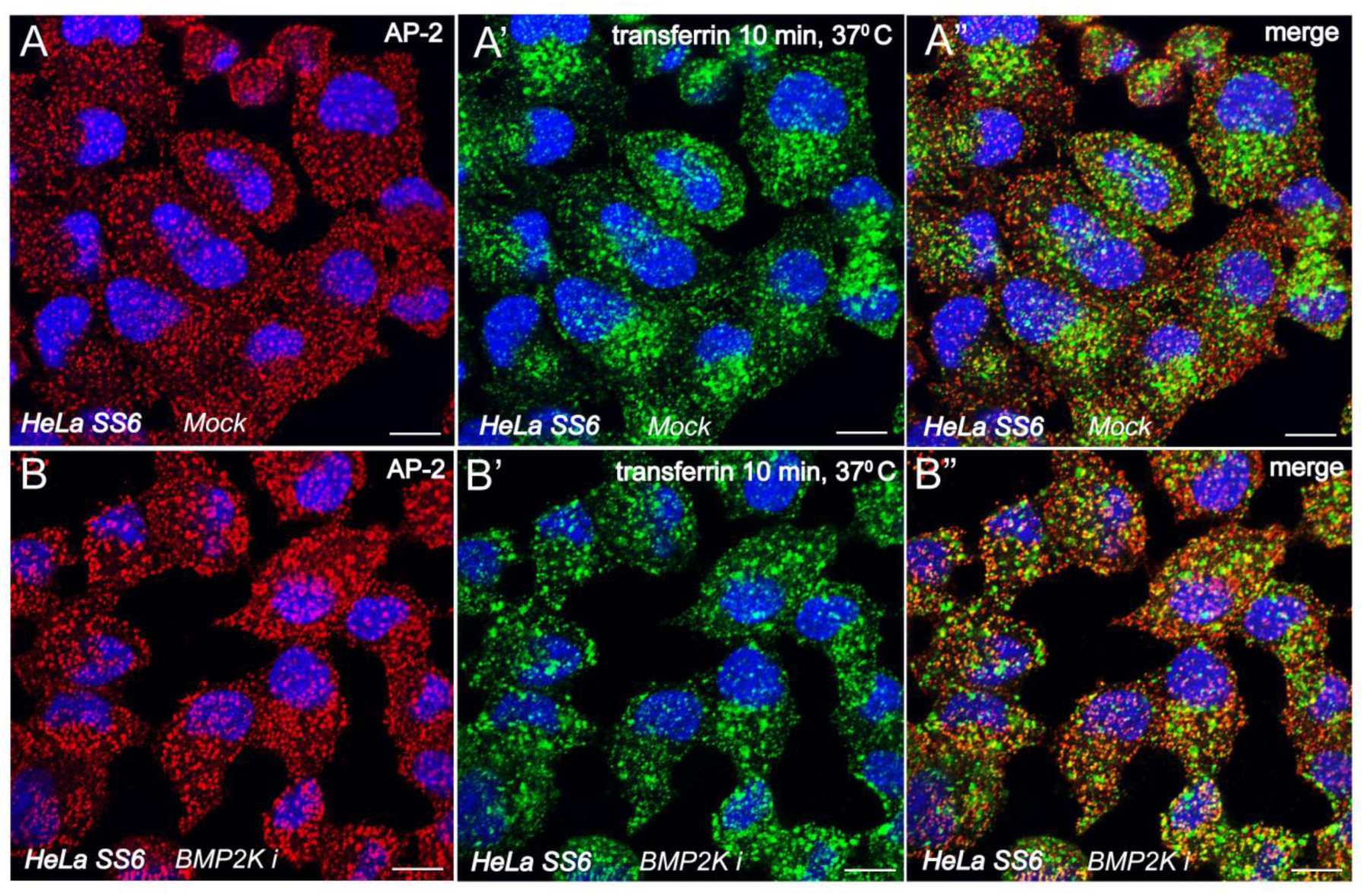
Persistent CCP maturation defects in BMP2K inhibited cells. (**A and B**) Representative confocal optical sections showing separated and merged colour channels of HeLaSS6 cells treated with 10 µM DMSO-Mock (**A**) or SGC-AAK1-1 inhibitor (BMP2Ki) (**B**) for 2 hours, pulsed with Alexafluor488-transferrin (green) for 10 minutes, stained for AP-2 (red) and DNA (blue). Scale bar-10 µm.

**Figure 9- figure supplement 1:**
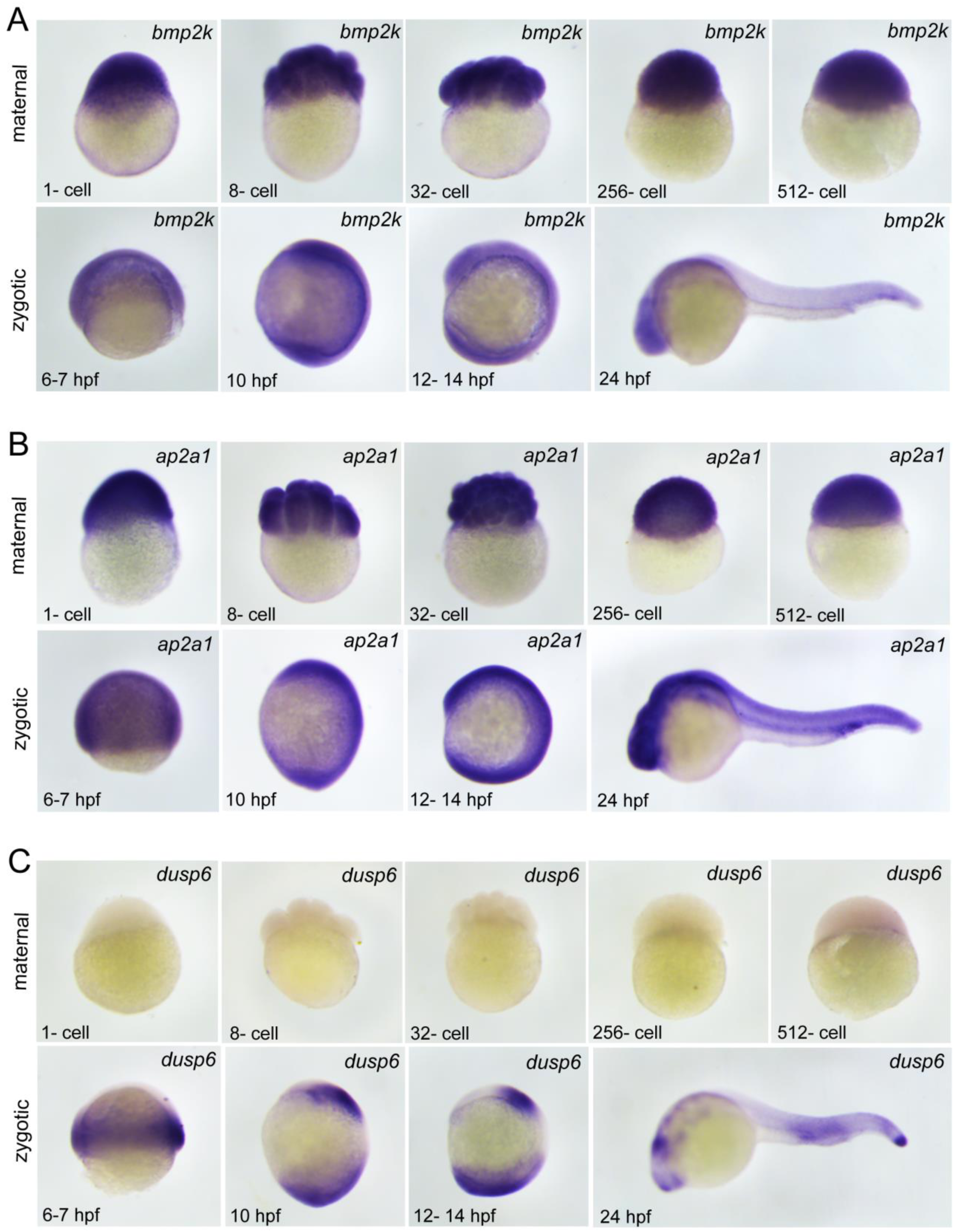
Analogous expression pattern of AP-2 and BMP2K during embryonic development. (**A-C**) Whole mount *in situ* analysis showing *bmp2k* (**A**), *ap2a1* (**B**) and *dusp6* (**C**) transcript localization in wild-type *D. rerio* embryos at the indicated maternal and zygotic developmental stages.

**Figure 9- figure supplement 2:**
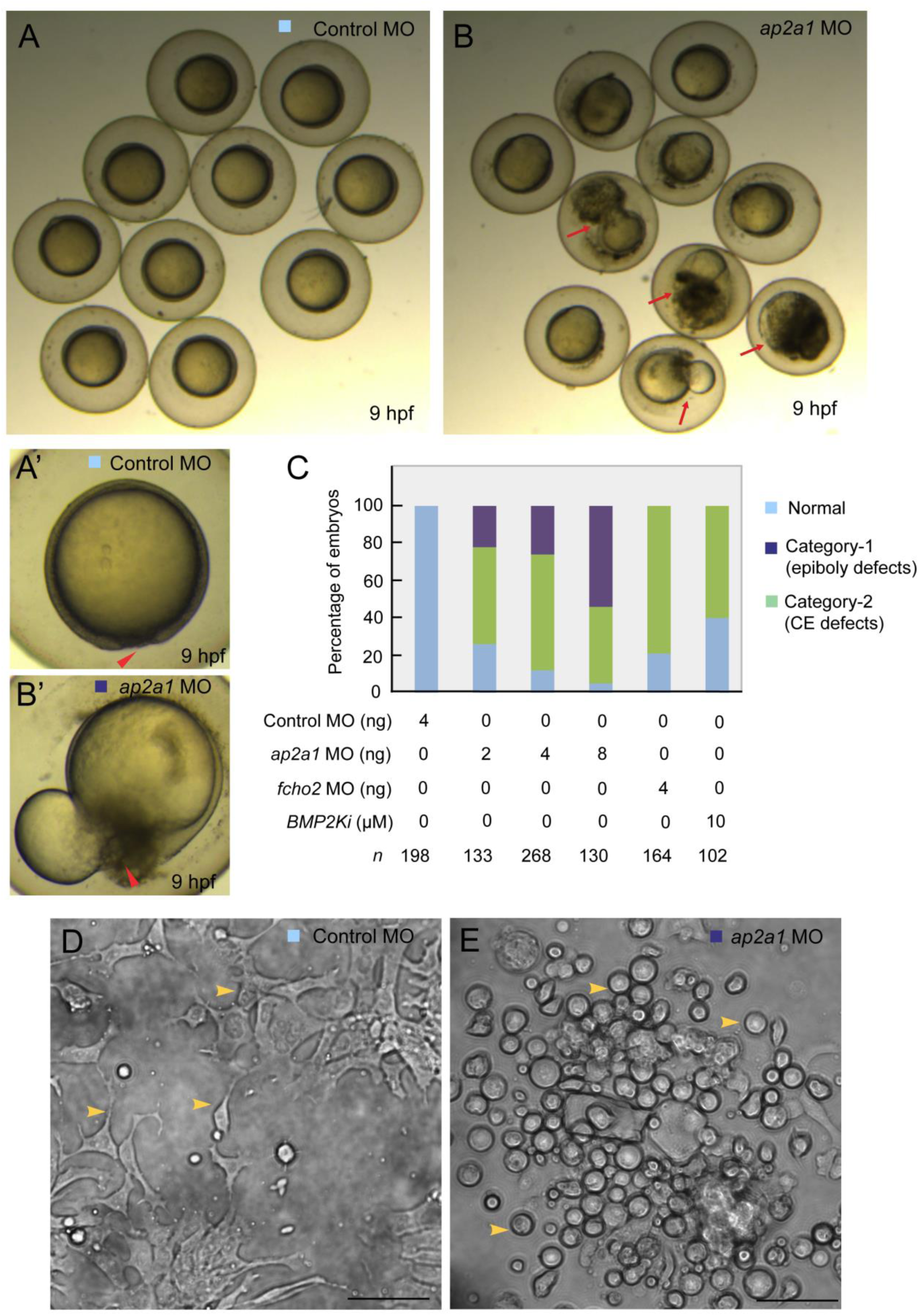
Embryonic defects in AP-2 morphants. (**A-B’**) Representative bright field live images showing comparative morphology of control (**A**) and *ap2a1* morphant (**B**) *D. rerio* embryos in clusters at 9 hpf. Dead or dying embryos (red arrows) during epiboly due to yolk extrusion through the blastopore (red arrow heads) are indicated. Zoomed images indicate morphology of typical normal (**A’**) and category-1 morphants (**B’**). Colour coded as in **C**. (**C**) Bar graphs showing quantification of normal (pale blue), and loss-of-function phenotypes belonging to category-1 (dark blue) that show epiboly defects and category-2 (pale green) that show convergent extension defects (CE) *n*: number of embryos analysed. (**D** and **E**) Representative phase-contrast images of cells isolated and cultured from epiboly stage *D. rerio* embryos injected with control (D) or *ap2a1* (E) morpholinos. Yellow arrow heads denote difference in cell morphology and adherence in control and *ap2a1* morphants. Scale bar-50 µm

